# Altered therapeutic capacities of olfactory ensheathing cells caused by a lesion in an autologous transplantation model for the treatment of spinal cord injury

**DOI:** 10.1101/2025.06.20.660789

**Authors:** Quentin Delarue, Matthis Brodier, Pauline Neveu, Laurine Moncomble, Clémence Raimond, Amandine Robac, Alizée Hugede, Axelle Blondin, Fannie Semprez, Pamela Lecras, Gaëtan Riou, Nicolas Guérout

**Affiliations:** Normandie univ, UNIROUEN, UR3830 GRHVN, Institute for Research and Innovation in Biomedicine (IRIB), 76000 Rouen, France; Université Paris Cité, CNRS UMR8003, Saints-Pères Paris Institute for the Neurosciences, F-75006 Paris, France; Normandie Univ, UNIROUEN, PRIMACEN Facility, IRIB, 76130 Mont-Saint Aignan, France & PBS-UMR6270 CNRS, FR3038 CNRS, 76130, Mont-Saint Aignan, France; Normandie univ, UNIROUEN, INSERM U1234, CHU Rouen, Department of Immunology and Biotherapy, F-76000 Rouen, France

## Abstract

Spinal cord injury (SCI) causes irreversible loss of motor, sensory, and autonomic functions and currently has no cure. Beyond local damage, SCI induces systemic inflammation, including cerebral inflammation that impairs neurogenesis. While cell therapies show promising effects in animal models, such as scar reduction and neuroprotection, their benefits in humans remain limited. One key difference lies in the transplantation strategy: animals receive healthy donor cells, whereas humans require autologous transplants. This led us to investigate how the lesion context affects the neuro-reparative potential of olfactory ensheathing cells (OECs) harvested from olfactory bulbs.

To this end, we cultured OECs from healthy animals and from animals that had undergone SCI one week earlier. We then transplanted both types of OECs into recipient animals after SCI for therapeutic purposes. Using functional sensory-motor studies, histological and gene expression analyses, we were able to demonstrate for the first time that the lesion negatively affects the therapeutic properties of cells used to treat SCI. Indeed, transplantation of cells from previously injured animals does not modulate the fibrotic and glial scar, or the demyelinated areas at the lesion site, and therefore fails to improve functional recovery; unlike cells derived from healthy donors. Moreover, our *in vitro* studies show that cells derived from SCI animals secrete pro-inflammatory molecules that promote the polarization of microglia toward a pro-inflammatory phenotype. Altogether, these innovative findings provide new insights into the potential of cell transplantation in the context of autologous therapy after SCI.

**Graphical abstract:** 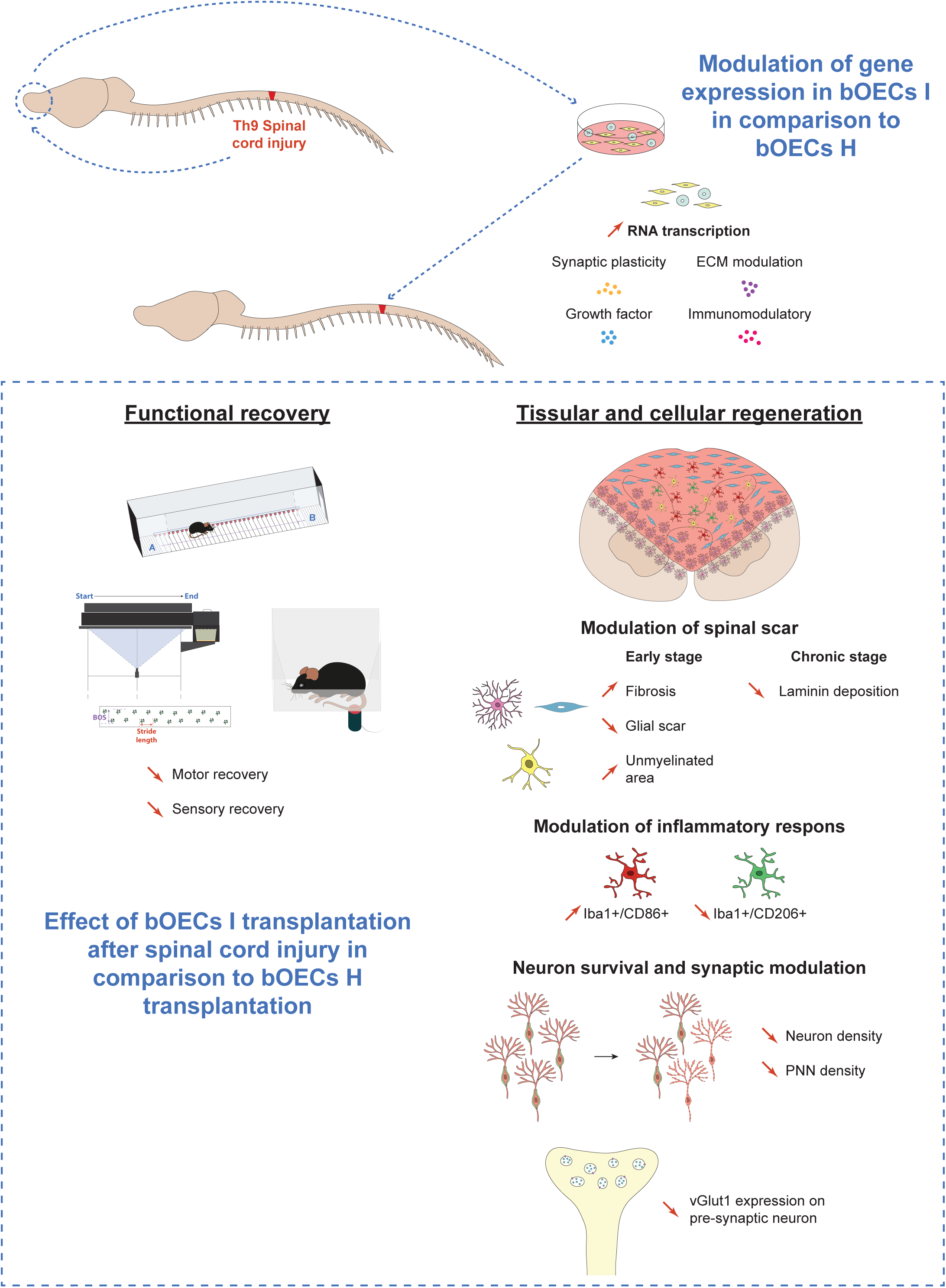

## INTRODUCTION

Traumatic spinal cord injury (SCI) is a relatively uncommon neurological condition most often caused by road-traffic, sports accidents, falls, or acts of violence (1). It is more prevalent among young individuals and those over 60 who are typically in good health (2). Unfortunately, no curative treatment is currently available for individuals with SCI, leaving them with lifelong disabilities. Depending on the severity of the injury, SCI can lead to complete or partial motor, sensory, and/or autonomic deficits below the injury level.

The pathophysiology of SCI is now better understood. Physical damage to the spinal cord, known as the primary lesion, triggers the activation of cellular and molecular mechanisms. Tissue destruction results in blood vessel rupture and cell death, which in turn leads to neuronal dysfunction. The release of cellular debris and cytokines into circulation leads to the infiltration and activation of neutrophils and macrophages, which then activate resident immune cells (microglial cells), triggering an inflammatory response (3,4). The emergence of inflammation leads to secondary damage, which is associated with the death of surrounding neurons. Moreover, this inflammation also activates astrocytes, ependymal cells, and pericytes, which contribute to the formation of a spinal scar. This scar helps to limit the spread of secondary damage and promotes spinal cord regeneration, albeit to a limited extent (5–7). The inflammatory response is not confined to the spinal cord. Indeed, it has been reported that SCI induces systemic and brain inflammation, particularly in the hippocampus and olfactory bulbs (OB), resulting in decreased neurogenesis and cognitive decline in mice and rats (8,9).

Numerous experimental and clinical approaches have been explored to provide a curative therapy for patients with SCI. Some of these strategies focus on conditioning the lesion environment using enzymatic digestion, hydrogels, and other biomaterials, to promote axonal regrowth. More recently, brain–machine interfaces have been engineered to bypass the damaged segment and re-establish communication between the brain and spinal networks distal to the lesion (10–12). Cell transplantation is among the most intensively studied interventions. Various cell types have been explored following SCI, including stem cells such as hematopoietic or mesenchymal stem cells (13,14), as well as differentiated cells such as Schwann cells (15) or olfactory ensheathing cells (OECs) (16). Despite their differing origins, all these cell types appear to converge on similar reparative mechanisms, most notably remodeling the glial–fibrotic scar, limiting secondary injury and inducing axonal regrowth/survival, which lead to improvement of locomotor functions (17).

Among the various cell types investigated, primary OB cultures, composed predominantly of bulbar olfactory ensheathing cells (bOECs) and fibroblasts, have emerged as a particularly promising candidate. In animal models of SCI, bOEC transplantation exerts both anti-inflammatory and pro-regenerative effects, improving functional recovery. Indeed, in their recently published manuscript, Phelps et al. highlight that the vast majority of studies have reported positive effects of OEC transplantation after SCI (18). Mechanistically, transplanted bOECs promote axonal survival and regrowth while re-shaping post-traumatic immunity, shifting macrophage–microglial (Mφ–Mi) from a pro-to an anti-inflammatory phenotype (19). These cells have progressed to clinical testing because they can promote tissue repair without permanently integrating into the spinal parenchyma. Human trials, however, have produced inconsistent outcomes: some report gains in American Spinal Injury Association (ASIA) scores, whereas others find little or no benefit (20). Divergent culture conditions, surgical techniques, and species-specific pathophysiology may account for these discrepancies. An additional difference is the transplantation paradigm itself: human studies rely on autologous grafts harvested from the injured patient, whereas most pre-clinical experiments use cells from healthy donors. Given that SCI elicits systemic and cerebral inflammation (9,21,22), we hypothesized that the injury environment could compromise the reparative capacity of cells collected post-trauma. Here, we test this hypothesis by characterizing bOEC transplantation harvested from mice after SCI and evaluating their therapeutic efficacy in an autologous-like transplantation model.

## MATERIALS AND METHODS

### Animal experimentation

#### Animal care and use statement

The experimental protocol was designed to minimize pain and discomfort for the animals. All experimental procedures adhered to the European Community guidelines on the care and use of animals (86/609/CEE; Official Journal of the European Communities no. L358; 18 December 1986), the French Decree no. 97/748 of 19 October 1987 (Journal Officiel de la République Française; 20 October 1987), and the recommendations of the Cenomexa Ethics Committee (#26905).

#### Animals

Mice of both sexes were housed in conventional, secure rodent facilities (two to five animals per cage, sexes separated) under a 12-h light/dark cycle with ad libitum access to food and water. A total of 270 adult mice (8–12 weeks old; mean body mass ≈ 20 g for females, 25 g for males) were used:

- Wild-type (WT) C57BL/6: 254 mice
- Rosa26-tdTomato: 12 mice
- Luciferase (LUX) line—CAG-luc-GFP (L2G85Chco+/+; FVB-Tg(CAG-luc,-GFP) L2G85Chco/J, JAX #008450): 4 mice, backcrossed to C57BL/6 for three generations to minimize strain-related variability

Each experimental group was balanced for sex (50 % male / 50 % female) and organized as follows (Figure 1B):

1. Non-injured control: unlesioned animals, providing baseline for functional analyses.
2. SCI: animals subjected to SCI without cell transplantation.
3. SCI + bOEC-H: SCI animals receiving bOECs cultured from healthy, uninjured donors, paradigm classically used in animal studies.
4. SCI + bOEC-I: SCI animals receiving bOECs cultured from donors that had themselves undergone SCI, modeling an autologous-transplant scenario.

**Figure 1:**
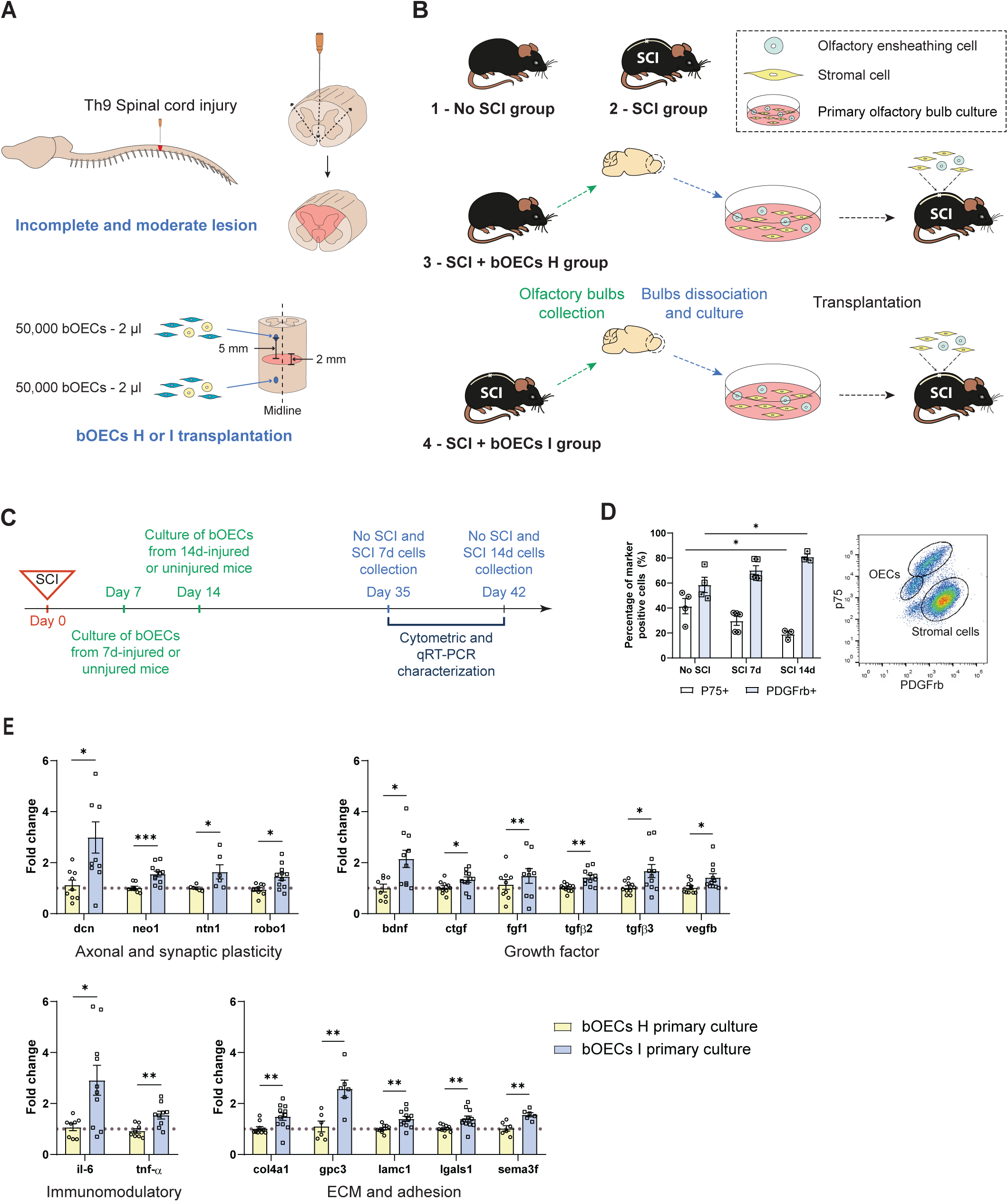
Experimental design and characterization of bOEC cultures. **A.** Schematic representation of Th9 spinal cord injury and bOECs transplantation. **B**. Schematic representation of the four main experimental groups used in our study. (1) No SCI group, uninjured animals used to define the baselines for functional experiments. (2) SCI group, injured animals without treatment. (3) SCI bOECs H, these animals underwent SCI and primary bOECs were transplanted immediately. OB from uninjured animals were used to prepare bOEC cultures. (4) SCI bOECs I, these animals underwent SCI and primary bOECs were transplanted immediately. OB from injured animals were used to prepare bOEC cultures. This SCI was performed on a different set of mice 7 days prior to the primary OB cultures. **C.** Experimental design, at day0, SCI was conducted. Healthy or injured bOECs were plated 7 and 14 days after SCI. Three weeks after plating, 35 days and 42 days, bOECs H and bOECs were collected and cytometric and qRT-PCR characterization was conducted. **D.** Quantification of the percentage of p75+ OECs and PDGFRβ+ fibroblasts in bOEC cultures without SCI, 7 days and 14 days after SCI by cytometric analyses. **E**. Histograms of bOECs H and bOECs I mRNA expression of immunomodulatory, axonal plasticity, ECM/signaling and growth factor genes. Dashed lines correspond to mRNA expression from bOEC H cultures used as control. Quantifications are expressed as average ± SD. N=4 (**D**) and 8 (**E**) animals per group. Statistical evaluations were based on Mann-Whitney tests. * = *P*< 0.05, ** = *P*< 0.01 and *** = *P*< 0.001.

#### Surgical procedure and cell transplantation

SCI was performed at the T9-T10 level as previously described (19). After a midline dorsal incision and laminectomy of the T7 vertebra, the cord was partially transected with a 25-gauge needle, sparing the ventral horn’s lower portion but fully severing the central gray matter and dorsal columns (Figure 1A). Immediately afterward, bOECs were transplanted with a micromanipulator-mounted stereotactic arm (World Precision Instruments). A sterile 1 mm glass capillary was lowered 1 mm deep, 1.5 mm lateral to the midline, 5 mm rostral and 5 mm caudal to the lesion. At each of the two sites, 2 µL of cell suspension (25 000 cells/µL) were injected over 1 min (Figure 1A). The musculature and skin were sutured and animals were monitored daily; none of the mice developed skin lesions, infections, or autotomy during the study.

#### Neuroanatomical tracing - Biotinylated dextran amine (BDA) injection

To trace axonal growth, 2 µL of biotinylated dextran amine (BDA; 0.2 g/mL, Thermo Fisher Scientific, D1956, Waltham, MA) diluted in PBS were injected at the cervical level 48 hours before euthanasia. Using a 1 mm sterile glass capillary mounted on a micromanipulator and stereotaxic frame (World Precision Instruments), we lowered the needle 1 mm deep and 1.5 mm lateral to the midline, then infused the tracer over 1 minute.

#### bOECs survival analysis

bOECs survival was assessed by two complementary methods. First, LUX+/− bOECs were prepared from OB of luciferase-reporter mice (see above for mouse line) and transplanted into wild-type recipients immediately after SCI. Bioluminescent emission from the grafted cells was captured with an *in-vivo* X-TREM 4XP cooled-CCD optical imager (Bruker). Mice received an intraperitoneal injection of D-luciferin (0.3 mg/g, XenoLight™, Perkin Elmer) and were imaged 30 minutes after injection, and then repeatedly up to three weeks after SCI. Photon counts were quantified with Bruker molecular-imaging software as previously described (23). To achieve greater sensitivity, we also transplanted bOECs isolated from Rosa26-tdTomato mice into wild-type recipients. The fluorescence of tdTomato-expressing bOECs was quantified by immunohistochemistry for up to four weeks post-transplantation (see below for histological analyses).

### Behavioral analyses

#### Locotronic foot misplacement test

Foot misplacement experiments (Intellibio, Nancy, France) was carried out as previously described by Chort et al. (24). The ladder links a start chamber to a goal box and is flanked by infrared sensors that record every step in real time. The dedicated software logs the position and duration of each misstep, discriminating among fore-limb, hind-limb, and tail errors. From these recordings we extracted two main outcomes, hind-limb error counts and total crossing time, for comparison across experimental groups (Figure 2A).

**Figure 2:**
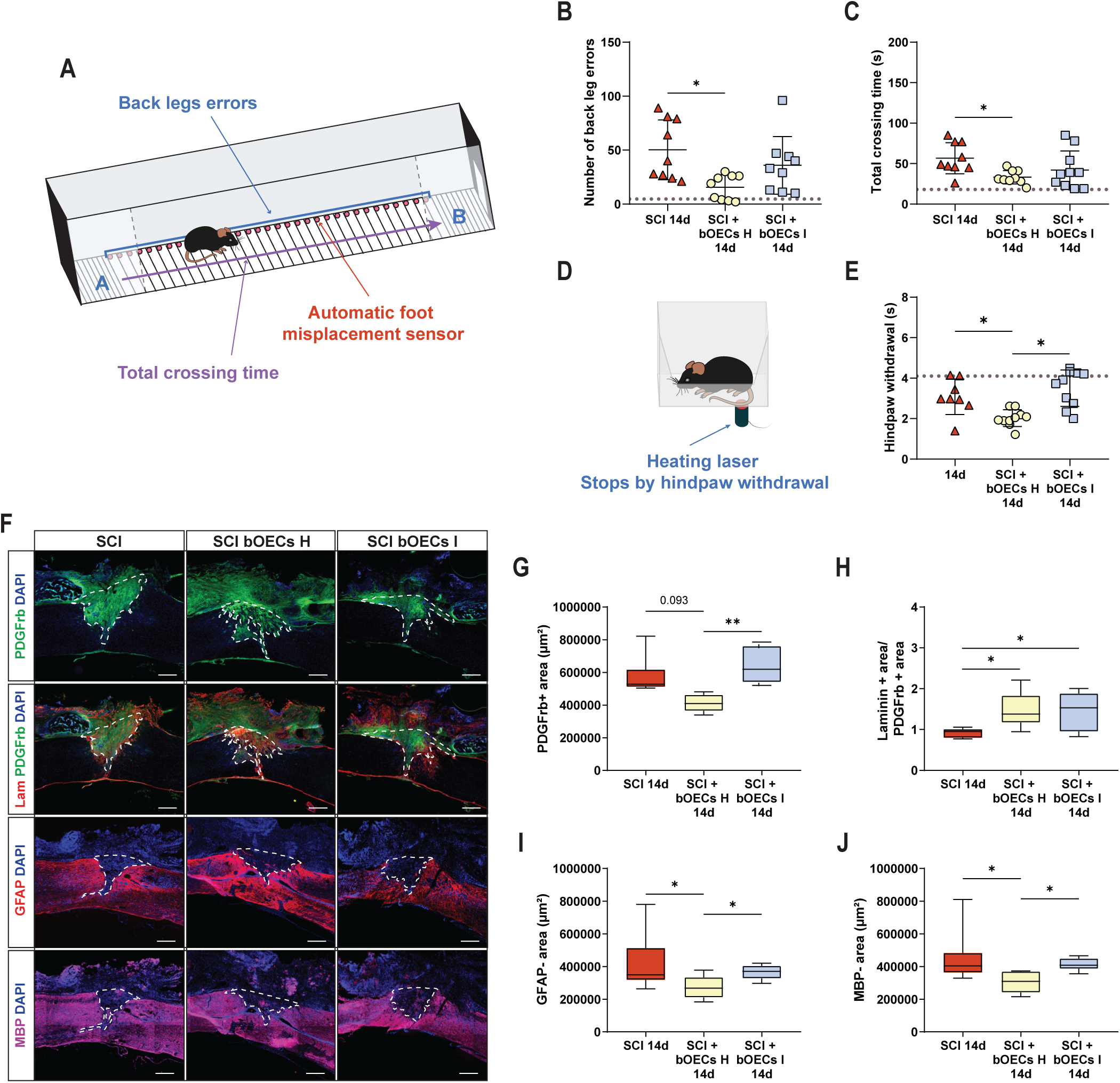
Transplantation of bOECs from injured mice resulted in less functional recovery and tissue regeneration than bOECs from healthy mice in early stage. **A.** Schematic representation of the Locotronic test. Mice traverse a corridor fitted with a horizontal ladder, and sensors positioned between the rungs automatically record locomotor errors as the animals move from point A to point B. **B–C.** Functional recovery was assessed with the Locotronic test 14 days after SCI. Dashed lines indicate baseline values recorded during habituation (7 days before SCI). **B.** Quantification of hindpaw errors 14d post-SCI. **C.** Quantification of total crossing time 14d post-SCI. **D.** Schematic representation of the Hargreaves test. A heating laser is applied to the plantar surface of each hindpaw, and the latency to paw withdrawal is measured. Greater sensitivity is reflected by a shorter withdrawal latency. **E.** Sensory recovery was analyzed using Hargreaves test 14d post-SCI. **F-J.** Histological analyses of the spinal scar were performed 14days after SCI. **F.** Representative pictures of sagittal spinal cord sections of SCI, SCI bOECs H and SCI bOECs I groups 14 days after SCI. Sections were stained with anti-PDGFRβ, anti-Laminin, anti-GFAP, and anti-MBP antibodies. **G.** Quantification of fibrosis areas (PDGFRβ+). **H.** Quantification of Laminin+ area/ PDGFRβ+ area ratio. **I.** Quantification of astrocyte-free areas (GFAP-). **J**. Quantification of demyelinated/unmyelinated area (MBP-). Dashed lines correspond to the injury site. Scale bar = 200 µm. N=8-10 (**B, C, E, G, H, I** and **J)** animals per group. Quantifications are expressed as average ± SD (**B, C** and **E**) and average ± Min/Max (**G, H, I** and **J)**. Statistical evaluations were based on Kruskal-Wallis tests. Quantifications are expressed as average ± SD. * = *P*< 0.05 and ** = *P*< 0.01.

All mice were pre-trained one week before injury. Behavioral testing was then performed at 14- and 56-days post-SCI.

#### Hargreaves plantar test

The Hargreaves plantar test (Ugo basile®) was conducted as previously described by Takano et al. (23). After a 5-min acclimation period, a focused radiant-heat source was applied to the plantar surface of each hind paw. The latency between stimulus onset and paw withdrawal, the hindpaw withdrawal latency, was recorded automatically (Figure 2D). Experiments were performed at 14- and 56-days post-SCI.

#### CatWalk XT automated gait analysis

Motor coordination was evaluated with the CatWalk XT automated gait-analysis system (Noldus; RRID:SCR_004074) (24). In a darkened room, each mouse crossed an illuminated glass walkway while a camera positioned beneath the platform captured the paw-print reflections. These images were digitized and processed with CatWalk XT 9.1, using an intensity threshold of 80 arbitrary units (range 0–225). After footprint detection and labeling, the software extracted static and dynamic gait parameters from three consecutive valid runs (Figure 3D). The average values of these parameters were then analyzed and compared across experimental groups. If a mouse paused or reversed direction, the trial was aborted and repeated until three uninterrupted crossings were obtained. All assessments were performed 56 days after SCI.

**Figure 3:**
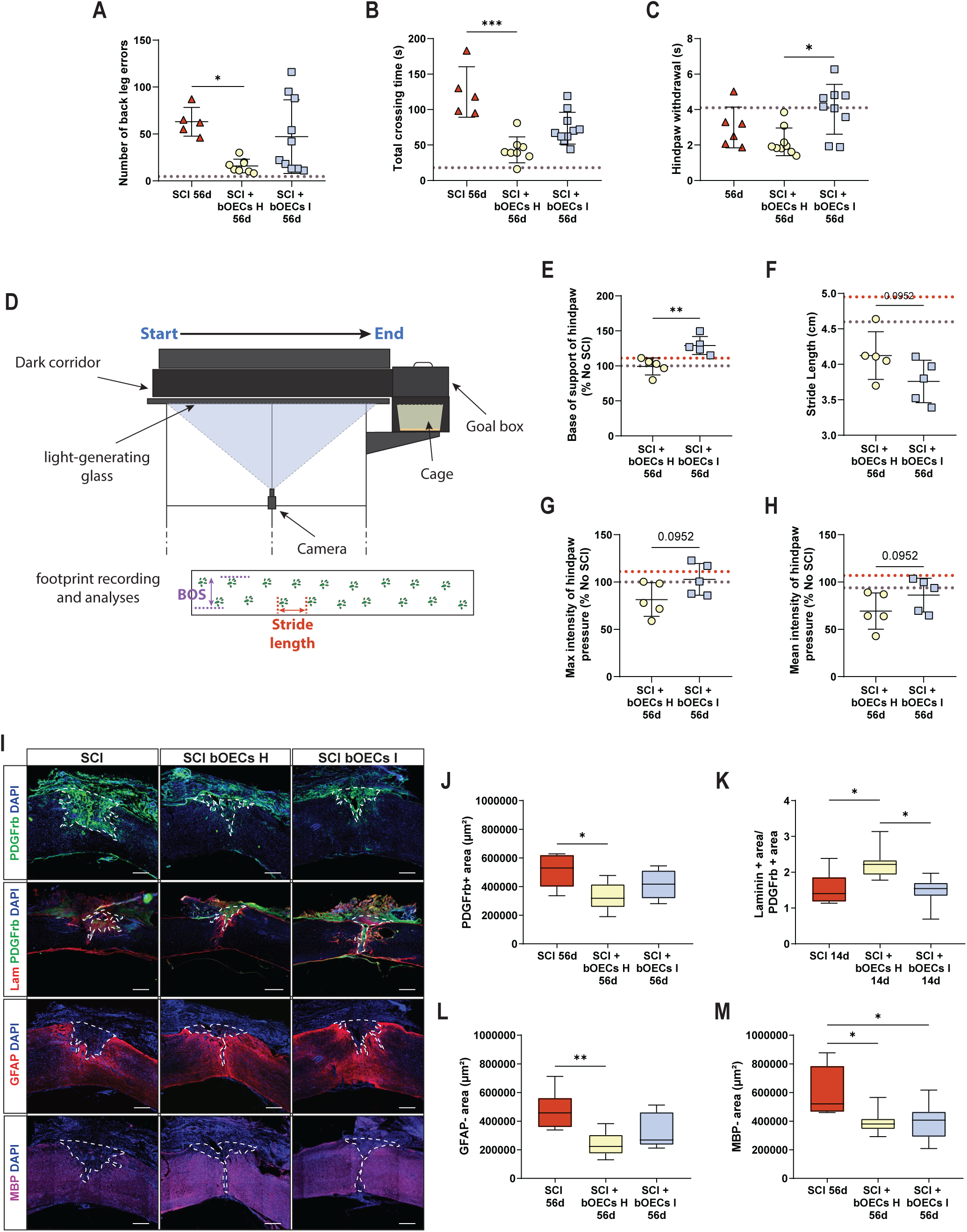
Modest therapeutic effects of bOEC I transplantation on functional and tissue regeneration after SCI in late stage. **A–B.** Functional recovery was assessed with the Locotronic test 56 days after SCI. Dashed lines indicate baseline values recorded during habituation (7 days before SCI). **A.** Quantification of hindpaw errors 56d post-SCI. **B.** Quantification of total crossing time 56d post-SCI. **C. S**ensory recovery was analyzed using Hargreaves test 56d post-SCI. **D-H.** CatWalk gait analysis was performed 56d after SCI. **D.** Schematic representation of the CatWalk test. Mice cross the dark corridor up to the goal box. Their footprints are then generated by the light-emitting glass and recorded by a camera. The distance between the hindpaws (BOS) and the stride length are measured. **E**. Measurement of the base of support (BOS) 56d post-SCI. **F**. Measurement of the stride length 56d post-SCI. **G.** Measurement of the max intensity of hindpaws 56d post-SCI. **H.** Measurement of the mean intensity of hindpaws 56d post-SCI. **I-M.** Histological analyses of the spinal scar were performed 56 days after SCI. **I.** Representative pictures of sagittal spinal cord sections of SCI, SCI bOECs H and SCI bOECs I groups 56 days after SCI. Sections were stained with anti-PDGFRβ, anti-Laminin, anti-GFAP, and anti-MBP antibodies. **J.** Quantification of fibrosis areas (PDGFRβ+). **K.** Quantification of Laminin+ area/ PDGFRβ+ area ratio. **L.** Quantification of astrocyte-free areas (GFAP-). **M**. Quantification of demyelinated/unmyelinated area (MBP-). Dashed lines correspond to the injury site. Scale bar = 200 µm. N=5-10 (**A, B, C, E, F, G, H, J, K, L** and **M)** animals per group. Quantifications are expressed as average ± SD (**A, B, C, E, F, G** and **H**) and average ± Min/Max (**J, K, L** and **M)**. Statistical evaluations were based on Kruskal-Wallis tests (**A, B, C, J, K, L** and **M**) and Mann-Whitney tests (**E, F, G,** and **H**). Quantifications are expressed as average ± SD. * = *P*< 0.05, ** = *P*< 0.01 and *** = *P*< 0.001.

### Cell transplant preparation and characterization

#### Primary OB culture and cell transplant preparation

Adult mice were anesthetized with 2 % isoflurane (Iso-Vet, Osalia, Paris, France) and euthanized by decapitation. OBs were rapidly removed, transferred to ice-cold phosphate-buffered saline (PBS), and incubated in 0.25 % trypsin (Thermo Fisher, 15090-046). After enzymatic digestion, the tissue was mechanically dissociated, and the resulting cell suspension was plated in T25 flasks containing 5 mL of DF-10S medium (DMEM-GlutaMAX supplemented with 10 % heat-inactivated fetal bovine serum and 0.5 % penicillin/streptomycin). Cultures were maintained until confluence (3–4 weeks), then trypsinized and counted. Finally, cells were resuspended at 25 000 cells/µL in DF-10S for transplantation.

#### Flow cytometry and bOECs characterization

Cell populations in bOEC cultures were analyzed by flow cytometry as described by Takano et al. (23). Cells were resuspended at 2×10^5^ cells/ml PBS/ 0.5% bovine serum albumin (BSA), and TruStain FcX™ PLUS (BioLegend) was added to block non-specific binding. Cell populations were identified using anti-p75 nerve growth factor receptor (p75, Abcam, ab8874, RRID:AB_306827) and rat anti-platelet-derived growth factor β (PDGFRβ, Abcam, ab91066, RRID:AB_10563302) primary antibodies. bOECs were identified as P75 positive (+) and PDGFRβ negative (-) cells. Stromal cells were identified as PDGFRβ+ and P75-cells. P75 and PDGFRβ primary antibody were detected with the anti-rabbit phycoerythrin fluorochrome-conjugated (PE, poly4064, BioLegend, 406408, RRID:AB_10643424) and the anti-rat Alexafluor 488 fluorochrome-conjugated (AF488, MRG2b-85, BioLegend, RRID:AB_2715913) secondary antibodies, respectively. Data were analyzed using FlowJo software (version 10.3; FlowJo LLC).

### BV2 cell culture in conditioned medium

#### bOEC conditioned-medium preparation

Conditioned medium (CM) was prepared from bOEC cultures derived from both uninjured and SCI mice. When cultures reached ∼80 % confluence, bOEC cultures were rinsed twice with PBS and then fresh medium was added. After four days, this medium was collected, transferred to Amicon Ultra-15 3 kDa centrifugal filters (Millipore), and spun at 5 000 × g for 1 h at room temperature. Concentration reduced the starting volume, from 6 mL, to ∼1 mL of CM, which was immediately aliquoted and stored at −80 °C until use.

#### BV2 Cell line cultures

The BV2 line is an immortalized murine microglial cell line (ATCC CRL-2467, EOC) derived from the brain of a 10-day-old female mouse. Vials were thawed in a 37 °C water bath, and the cells were transferred to T25 flasks containing high-glucose DMEM (Gibco) supplemented with 20 % fetal bovine serum (FBS) and 0.5 % penicillin/streptomycin. After 48 h, the cultures were first passaged: the medium was removed, cells were rinsed twice with PBS, exposed to 5 mL trypsin–EDTA for 10 min at 37 °C, and the reaction was quenched with complete medium. The suspension was centrifuged at 500 × g for 5 min, and the pellet was resuspended in 1 mL fresh medium. Routine culture was then carried out in high-glucose DMEM GlutaMAX containing 10 % FBS and 0.5 % penicillin/streptomycin (DF-10S high glucose). One week later, cells were passaged again and distributed into three experimental conditions (Figure 5I):

- 1-BV2 control: The medium consists of DF-10S high glucose
- 2-BV2 + bOECs H CM: The medium is composed of DF-10S high glucose supplemented with bOECs H CM
- 3-BV2 + bOECs I CM: The medium is formulated with of DF-10S high glucose and supplemented with bOECs I CM

After three days in culture, the cells were harvested by trypsinization, pelleted, and snap-frozen at −80 °C until qRT-PCR analysis.

#### Real-time quantitative polymerase chain reaction characterization

Quantitative reverse transcription polymerase chain reaction (qRT-PCR) experiments were conducted to measure the mRNA expression levels of axon growth inhibitory and permissive molecules, as well as neurotrophic factors in the bOEC cultures (Table S1). This first genes panel is based on the one published by Anderson et al. (5). Additionally, qRT-PCR was performed to evaluate the mRNA expression levels of cytokines, chemokines, and inflammatory molecules in BV2 cell cultures, including BV2 control, BV2 + bOECs H CM, and BV2 + bOECs I CM (Table S2). This second genes panel is based on the one used by Briffault et al. (25). Total RNA was extracted from bOECs and BV2 cells using Tri-reagent (Sigma) and the Nucleospin RNA II Kit (Macherey-Nagel) according to the manufacturer’s instructions. From each sample, 1.5 µg of total RNA was converted into single-stranded cDNA using the ImProm-II reverse transcriptase kit (Promega) with random primers (0.5 µg/mL). Two ng of complementary DNA (cDNA) were amplified in the presence of 2X SYBR Green Mastermix (Applied Biosystems), which contains preset concentrations of dNTPs, MgCl_2_, and both forward and reverse primers, using a QuantStudio™ 12K Flex (Applied Biosystems). Appropriate controls, including reactions without reverse transcriptase and those using H_2_O instead of cDNA, were included. Primers were designed using Primer Express software (Applied Biosystems) and validated through RT-PCR. The relative amount of cDNA in each sample was quantified using the comparative quantification cycle (Cq) method and expressed as 2⁻ΔΔCq. Three housekeeping genes *gapdh*, *b2m*, and *actb*, were employed to standardize the relative amount of cDNA. All experiments were performed with two technical replicas.

### Histological analyses

#### Tissue preparation and sectioning

Thirty minutes before euthanasia, mice received analgesia (buprenorphine, 0.17 mg/kg). They were then deeply anesthetized with 5 % isoflurane and given an intraperitoneal injection of sodium pentobarbital (Euthoxine, 100 mg/kg). Transcardial perfusion was performed with ice-cold PBS followed by ice-cold 4 % paraformaldehyde (PFA; Sigma-Aldrich 252549) in PBS. Spinal cords were dissected, post-fixed overnight in 4 % PFA, and cryoprotected for ≥ 48 h in 30 % sucrose (Life Technologies). Tissues were embedded in Tissue-Tek OCT compound (Sakura) and sagittally sectioned at 20 µm. Sections were mounted on slides and stored at −20°C until use.

#### Immunohistochemistry

Identification of different cells and structures in spinal cord were conducted using primary antibodies diluted in a PBS blocking solution containing 10% normal donkey serum (Jackson ImmunoResearch, Cambridge, UK) and 0.3% Triton-X100 (Sigma-Aldrich) to minimize non-specific binding. Antibodies incubation occurred overnight at room temperature in a humidified chamber. The following primary antibodies were used: rabbit anti-platelet-derived growth factor β (PDGFRβ, Abcam, ab32570, RRID:AB_777165), mouse anti-glial fibrillary acidic protein (GFAP Cy3-conjugated Sigma-Aldrich, C9205, RRID:AB_476889), rabbit anti-ionized calcium-binding adapter molecule 1 (Iba1, Wako, 019-19741, Osaka, Japan, RRID: AB_839504), rat anti-myelin basic protein (MBP, Millipore, MAB386, RRID:AB_94975), rabbit anti-laminin (Abcam, ab11575, RRID: AB_298179), goat anti-mMMR/CD206 (CD206, R&D systems, AF2535, RRID:AB_2063012), rat anti-B-lymphocyte activation antigen B7-2/CD86 (CD86, Invitrogen, MA1-10299, RRID:AB_11152324), guinea pig anti-vesicular glutamate transporter 1 (vGlut1, Millipore, AB5905, RRID: AB_2301751), mouse anti-RNA binding Fox-1 homolog 3 (NeuN, Sigma, HPA030790, RRID) and Wisteria floribunda agglutinin biotin-conjugated lectin (WFA, Sigma, L1516, RRID: AB_2620171).

After washing, antibody staining was detected with species-specific fluorescence-conjugated secondary antibodies (Jackson ImmunoResearch). WFA antibody and BDA staining were visualized with Cy3-conjugated streptavidin (Jackson ImmunoResearch). Sections were counterstained with 4′,6-diamidino-2-phénylindole (DAPI; 1 μg/mL; Sigma-Aldrich) and mounted with Vectashield mounting medium (Vector Labs, Burlingame, UK).

#### Image acquisition analysis and quantification

Representative images for quantifying immunohistochemically stained areas in sagittal sections were captured using the Zeiss Apotome2 microscope and the Leica THUNDER Imager Tissue 3D microscope. Images of the lesion epicenter were obtained using a 10x microscope objective. Image processing and assembly were conducted with ImageJ software.

Three to five sections per animal were examined to identify the epicenter of the lesion, and an image of this epicenter was subsequently captured and analyzed. In these sagittal sections, area GFAP-, PDGFrβ+, MBP-and Laminin+ were measured. Additionally, NeuN+, WFA+, Iba1+, Iba1+/CD86+, Iba1+/CD206+ cells and BDA+/vGlut1+ cells were counted after standardizing the area using a rectangle of 1000 µm x 1000 µm.

### Statistical analysis

All data are presented as mean ± standard deviation (SD). Statistical analyses were conducted using GraphPad Prism software, version for windows (GraphPad Software; 8.0.1.244).

For figures comparing two groups, the averages were assessed using the two-way Mann-Whitney test. For figures comparing three groups, median comparisons were performed using the two-way Kruskal-Wallis test, followed by Dunn’s post-test. Detailed statistical analyses for each assay are provided in the figure legends. A p-value of less than 0.05 was considered statistically significant.

## RESULTS

### SCI alters cell proportion and gene expression in bOECs cultures

To determine how SCI alters the cells used for transplantation, we quantified the cellular composition and gene-expression profile of bOEC cultures derived from OBs of uninjured mice (bOECs H) and from mice 7 or 14 days after SCI (bOECS I) (Figure 1C-E). Flow-cytometry analysis with anti-p75 (OEC marker) and anti-PDGFRβ (stromal-cell marker) antibodies revealed that cultures from uninjured OBs consisted of 41.5 ± 12.2 % of OECs and 58.5 ± 12.2% of stromal cells, consistent with previous reports (19,20). Seven days after SCI, the composition of the primary cultures was only modestly shifted (29.7 ± 8.1 % OECs; 70.3 ± 8.1% stromal), whereas by 14 days the relative proportions had changed significantly, with OECs falling to 19.2 ± 3.4 % and stromal cells rising to 80.8 ± 4.0 % (Figure 1D and Figure S1E). In cultures from uninjured mice, OECs make up between one-third and one-half of the cells, whereas their relative proportion drops to below one-third in cultures prepared 7 days after SCI and to about one-fifth in cultures prepared 14 days after SCI. Because cultures prepared 7 days post-injury still contained about one-third OECs, comparable to uninjured controls, we limited subsequent experiments to this time point to minimize any bias introduced by large shifts in cell proportions.

Then, to observe intrinsic transcriptional changes, we performed qRT-PCR on 7-day SCI-derived cultures and uninjured controls. Results show that SCI significantly modulates genes linked to extracellular matrix (ECM) remodeling and axonal plasticity processes, such as *sema3f*, *neo1*, *ntn1* or *lgals1*. In addition, SCI changes the expression of immunomodulatory (*il6* and *tnf-a*) and growth factor genes (e.g. *bdnf* or *vegfb*) and genes involved in scar-forming molecules such as *ctgf*, *fgf1*, *tgfβ2* or *tgfβ3* (Figure 1E).

Collectively, our data indicate that SCI alters both the cellular composition and transcriptomic profile of bOECs, selectively upregulating genes involved in spinal cord tissue remodeling, cell survival, and immune regulation.

### The survival of bOECs derived from healthy mice and from SCI mice is identical

We questioned whether changes in gene expression could influence the survival of bOECs after transplantation and, consequently, impact their therapeutic efficacy. To address this question, we measured cell survival using bioluminescence and immunohistological assays.

Bioluminescence experiments enabled longitudinal analysis up to 21 days post-SCI in the same mice. For this purpose, bOECs H and I cultures were obtained from luciferase-expressing mice (bOECs Lux +/−). When exposed to luciferin, these cells emit photons that are captured by a bioluminescence imaging device. The results indicate that no surviving cells were detected in either transplanted group (bOECs H or bOECs I) 21 days post-SCI (Figure S1A and B). Additionally, these experiments revealed no significant differences between the two groups 7 and 14 days after SCI (Figure S1A and B).

The bioluminescence assay has a detection threshold that prevents the identification of a small number of cells. To circumvent this issue, we transplanted bOECs derived from tdTomato-positive mice into wild-type mice and conducted immunohistological analyses at 14, 21, and 28 days post-SCI. Although this technique is cross-sectional, it is more sensitive than bioluminescence. Measurements of tdTomato intensity showed no difference between the two groups at 14 and 21 days after SCI, confirming the bioluminescence results. Moreover, this experiment provided more precise measurements of bOECs survival and showed that, at 28 days, a small number of bOECs, identical in both groups, was still detectable (Figure S1C and D). Consequently, we observed no influence of the lesion on the survival of bOECs obtained from SCI mice after transplantation.

### bOECs I transplantation does not promote tissue regeneration and functional recovery during the early stage after SCI

To determine whether changes in RNA expression in cultured bOECs influence their neuroregenerative capacity after transplantation in the context of SCI, we evaluated functional and tissue recovery at an early stage 14 days post-SCI. Functional recovery was assessed with the Locotronic foot-misplacement test (Figure 2A–C) and the Hargreaves plantar test (Figure 2D and E), which evaluate motor and sensory function, respectively.

Analysis of total crossing time and hindpaw errors showed that transplantation of bOECs H, compared with non-transplanted mice, improved functional recovery, as evidenced by faster corridor crossing and fewer hindpaw misplacements. In contrast, transplantation of bOECs I did not result in significant improvement compared to non-transplanted mice. Furthermore, no significant differences were observed between the bOECs H and bOECs I groups, suggesting that bOECs I possess a lower therapeutic potential to promote motor recovery. (Figure 2A–C). It has been previously reported that cell transplantation, including OEC transplantation, can induce allodynia after SCI, reflecting a sensory-recovery phenomenon (26). To investigate this, plantar sensitivity was evaluated with the Hargreaves test 14 days post-SCI. In this test, a heating laser was applied to the sole of the mice’s hindpaw, and the latency to paw withdrawal, marking the onset of painful sensation, was recorded. A shorter withdrawal latency corresponds to greater sensory recovery. Mice transplanted with bOECs H show hindpaw-withdrawal latency was reduced compared with both mice transplanted with bOECs I and non-transplanted mice which displayed identical values (Figure 2D and E). Together, these two tests demonstrate that, relative to bOECs I transplantation, bOECs H transplantation promotes early sensory-motor recovery after SCI.

To correlate these functional data with tissue recovery, spinal-cord immunohistological analyses were performed 14 days post-SCI. We measured the fibrotic and astrocytic components of the lesion scar. The size of the fibrotic scar is a key factor in tissue and axonal regeneration after SCI: it inhibits regrowth by acting as a physical barrier. Conversely, the glial scar surrounding the fibrotic core expresses permissive factors for axonal regrowth. As the fibrotic scar enlarges, the astrocyte-free zone increases. Accordingly, we characterized the glial scar by quantifying the GFAP-area and assessed demyelination by measuring the MBP-area. As previously reported, transplantation of bOECs H modulated the lesion environment by reducing the fibrotic scar (PDGFRβ+ area) (Figure 2F and G), enlarging the glial component of the scar (reduction of GFAP-area) (Figure 2F and I), and thereby diminishing the unmyelinated region (Figure 2F and J) at 14 days post-SCI compared with the SCI control group. In contrast, transplantation of bOECs I did not induce comparable scar modulation (Figure 2F, G, I and J). Indeed, the size of the spinal-cord scar in mice transplanted with bOECs I is comparable to that of non-transplanted animals (Figure 2F, G, I and J).

The ECM composition plays a crucial role in both axonal regrowth and inhibitory processes. Accordingly, we examined the distribution of laminin (Figure 2F and H), an ECM molecule that promotes axonal growth. Because the laminin+ area is proportional to the size of the lesion core (PDGFRβ+ area), we calculated the ratio between the laminin+ and PDGFRβ+ areas. Our data show that, 14 days post-injury, this ratio is increased in both transplanted groups relative to the SCI control, with no significant difference between the two transplanted groups in terms of laminin deposition.

Taken together, these findings demonstrate that, at the acute stage, bOECs H confer a greater therapeutic benefit than bOECs I, leading to superior functional recovery and tissue repair.

### bOEC I transplantation exerts only modest therapeutic effects on both functional recovery and tissue regeneration in the context of late-stage spinal cord injury

We resumed the comparison of bOECs H and bOECs I transplantation at a later stage by transplanting a new cohort of animals and repeating the same analyses 56 days post-SCI. We also performed CatWalk analysis, a dynamic locomotor test that quantifies gait quality. The apparatus consists of an internally illuminated glass walkway that captures paw prints as the animals walk, recorded by a high-speed camera. CatWalk is more sensitive than the Locotronic hindpaw misplacement test and provides a complementary, detailed assessment of locomotion (Figure 3D-H).

At this chronic stage of SCI pathology, foot misplacement analysis shows that transplantation of bOECs I has little to no therapeutic effect (Figure 3A and B). Indeed, bOECs I transplantation did not differ significantly from either the SCI control or the bOECs H group in these parameters. In contrast, bOECs H transplantation yielded significant functional improvement 56 days post-SCI, as evidenced by fewer errors (Figure 3A) and shorter crossing times (Figure 3B) in comparison to control SCI mice. In addition to these motor analyses, comparisons of the base of support (BOS) between bOECs H and bOECs I groups was conducted. It appears that the distance between the hindpaws, and stride length further was performed and showed that bOECs H transplantation improved gait quality, reflected by a reduced distance between the hindpaws (Figure 3E). Although not statistically significant, the bOECs I group displayed a downward trend in stride length (p = 0.0952), suggesting a smaller functional improvement than that achieved with bOECs H (3.8 ± 0.3 cm vs 4.1 ± 0.3 cm) (Figure 3F).

Plantar-sensitivity analysis was also conducted at this late stage and yielded results similar to those at the early stage; mice transplanted with bOECs H withdrew the hind paw more quickly than those transplanted with bOECs I (Figure 3C). Moreover, the heightened sensitivity in bOEC-transplanted mice was partly echoed in the CatWalk test, which measures hindpaw pressure. Although the differences were not statistically significant, a downward trend was apparent in both mean (69.3.5 ± 19.1 86.2 ± 17.9 %) and peak (81.5 ± 17.8 % vs 102.8 ± 16.8%) pressure in bOECs H–treated mice relative to the bOECs I group (Figure 3G and H). Immunohistological analyses of the spinal cords of bOECs I-transplanted mice 56 days post-SCI revealed modest improved tissue regeneration (Figure 3I-M). Indeed, and as previously observed, transplantation of bOECs H induced a significant modulation of the astrocytic and fibrotic components of the spinal cord scar. Similarly, transplantation of bOECs I also led to modulation of these two parameters, although this effect was not statistically different from that observed in control animals (Figure 3J and L). Moreover, measurement of the MBP-area showed identical effects of bOECs H and bOECs I transplantation at 56 days post-SCI; both treatments reduced the unmyelinated area compared with the SCI group (Figure 3M).

In contrast to the early stage, the laminin composition of the spinal scar differs between the bOECs H and bOECs I groups 56 days after SCI. In effect, quantification of the laminin+ to PDGFRβ+ area ratio shows that bOECs H transplantation increases the relative amount of laminin within the scar compared with both the bOECs I-transplanted and non-transplanted groups (Figure 3K). This suggests that, at this late stage, the scar environment is more permissive for axonal growth after bOECs H transplantation.

These late-stage results underscore the beneficial effect of bOEC transplantation on both functional recovery and tissue regeneration after SCI. However, the magnitude of the benefit differs: bOECs I provide only a modest therapeutic effect compared with the more substantial improvements achieved with bOECs H.

### Transplantation of bOECs I fails to promote neuronal survival and synaptogenesis

Previous studies have demonstrated the ability of bOECs to enhance neuronal survival, neurogenesis, and axonal regrowth (18,27). Thus, we assessed neuronal density by immunohistochemistry using an anti-NeuN antibody to identify neurons. Neuronal density was quantified in the no-SCI group and in the three SCI groups at 14, 28, and 56 days post-SCI (Figure 4A–D). NeuN+ cells were counted in the rostral, caudal, and lesion-core parts of the spinal cord. Our results show that, 14 days post-SCI, neuronal density was increased only in bOECs H-transplanted mice, and only in the rostral part of the spinal cord, relative to the SCI group (Figure 4B). By 28 days, bOECs H transplantation had increased the number of NeuN+ cells in both the rostral and caudal regions compared with the SCI group (Figure 4C). Finally, at 56 days post-SCI, no significant differences were detected among the SCI groups (Figure 4D). These findings demonstrate that, unlike bOEC H transplantation, bOEC I transplantation does not improve neuronal density at 14, 28, or 56 days post-SCI.

**Figure 4:**
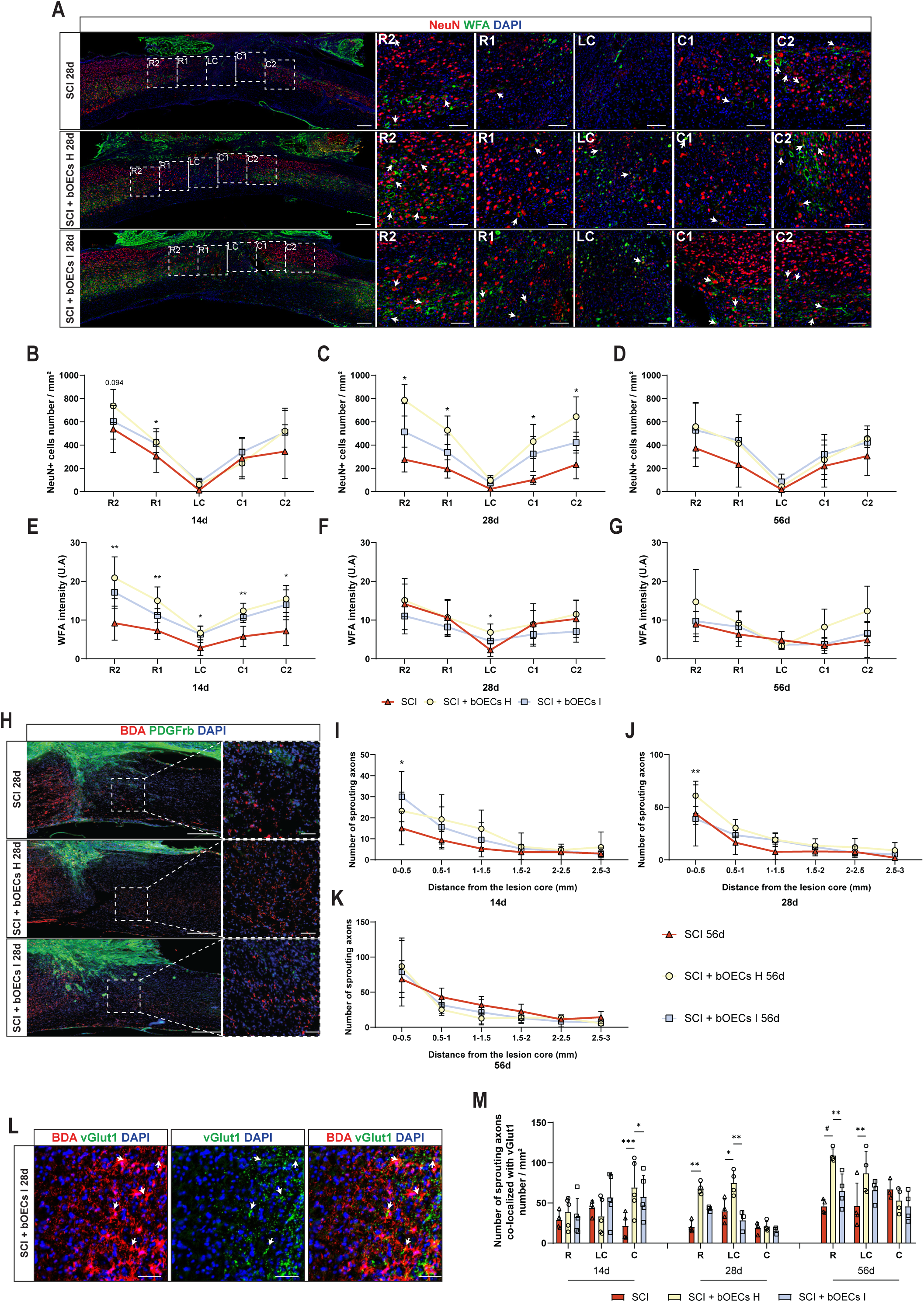
Transplantation of bOECs from injured mice fails to promote neuronal and synaptic preservation like bOECs H transplantation. **A**. Representative images of sagittal spinal cord sections from SCI, SCI bOECs H, SCI bOECs I groups 28 days after SCI. Sections were stained with anti-NeuN antibody and biotinylated WFA staining. The square sections correspond to the rostral 1 and 2 (R1 and R2), lesion core (LC) and caudal 1 and 2 (C1 and C2) areas, defined for the quantification of NeuN+ neurons and WFA+ perineural net. Arrows indicate co-localized NeuN+/WFA+ neurons. Scale bar = 200 µm and 100 µm. **B-D.** NeuN+ cell quantification in three experimental groups at 14d (**B**), 28d (**C**) and 56d (**D**). **E-G.** WFA intensity measure in three experimental groups at 14d (**E**), 28d (**F**) and 56d (**G**). **H.** Representative images of sagittal spinal cord sections from SCI, SCI bOECs H, SCI bOECs I groups 28 days after SCI. Sections were stained with anti-PDGFrβ antibody and biotinylated BDA staining. Scale bar = 200 µm and 100 µm. **I-K.** The number of sprouting axons as a function of distance from the lesion core was counted 14, 28 and 56 days after SCI. **L.** Representative images of sagittal spinal cord sections from SCI bOECs I groups 28 days after SCI. Sections were stained with anti-vGlut1 antibody and biotinylated BDA staining. Arrows indicate co-localized BDA+/vGlut1+ neurons. Scale bar = 50 µm. **M.** Sprouting axons co-localized with synaptic vGlut1+ were counted 14, 28 and 56 days after SCI. Quantifications are expressed as average ± SD (**B, C, D, E, F, G, I, J, K,** and **M**). 8-10 (**B, C, D, E, F** and **G**) and 4-5 (**I, J, K,** and **M**) animals per group. Statistical evaluations were based on Kruskal-Wallis tests. * = *P*< 0.05; ** = *P*< 0.01 and *** = *P*< 0.001.

To complete this neuronal analysis, we examined the perineuronal net (PNN) using WFA staining. This ECM structure is implicated in neuronal survival and synaptic plasticity; increased PNN density contributes to neuronal protection and synapse stabilization (28). Using immunohistochemistry, we quantified WFA intensity 14, 28, and 56 days post-SCI in the rostral, caudal, and lesion-core regions of the spinal cord (Figure 4A and E–G). Fourteen days after SCI, only bOEC H transplantation significantly increased WFA intensity in all three regions, whereas bOEC I transplantation showed a non-significant upward trend compared with the SCI group (Figure 4E). Analysis at 28 days revealed that bOEC H transplantation increased PNN density only within the lesion core (Figure 4F). By 56 days, no differences were detected among the groups. These data highlight distinct PNN dynamics across the three groups: early preservation is greater in bOEC H-treated mice than in SCI or bOEC I-treated mice, but from 28 days onward, except in the lesion core of the bOEC H group, PNN density is similar across groups. We can hypothesize that preservation of the PNN in bOEC H-transplanted animals supports neuronal survival at 14 and 28-days post-injury, whereas its reduction at 28 days preceded the decline in neuronal density observed at 56 days (Figure 4B-D).

As noted earlier, transplantation of bOECs I after SCI does not modulate spinal-scar size or the ECM in the same way as transplantation of bOECs H. The scar’s size and its ECM composition are critical determinants of axonal regrowth, inhibitory signaling, and synaptogenesis (29). We therefore conducted immunohistochemical experiments to characterize axonal growth and synaptogenesis, using the anterograde tracer BDA and the presynaptic marker vGlut1 (Figure 4H–M). As previously described (30), quantification of BDA+ axons within the lesion core showed that bOEC transplantation induced modest axonal regrowth after SCI compared with the SCI control. Specifically, mice transplanted with bOECs I displayed an increase in BDA+ fibers in the lesion core 14 days after injury, whereas bOECs H transplantation produced a similar increase at 28 days post-SCI (Figure 4I and J). By 56 days post-SCI, no significant difference was detected among the groups (Figure 4K). For both bOECs H and bOECs I, the increase in BDA+ fibers was confined to the lesion core; beyond 0.5 mm from the lesion, no differences were observed among groups at 14, 28, or 56 days post-SCI (Figure 4I–K).

In addition, colocalization of BDA and vGlut1 staining was analyzed 14, 28, and 56 days post-SCI (Figure 4L and M). The results showed that bOEC H transplantation increases the number of BDA+/vGlut1+ synapses in both the lesion core and in the rostral part of the spinal cord, compared with the SCI group, at 28 and 56 days. Relative to bOEC I transplantation, bOEC H also increased the number of BDA+/vGlut1+ synapses in the lesion core at 28 days and in the rostral segment at 56 days. A significant difference between the SCI and bOEC I groups was observed only in the caudal segment at 14 days, where bOEC I transplantation increased the number of BDA+/vGlut1+ synapses (Figure 4M).

Altogether, these results highlight that transplantation of bOECS H promotes neuronal survival, PNN expression, and synaptogenesis, whereas transplantation of bOECs I has little to no effect on these parameters.

### bOEC I transplantation promotes macrophage and microglial reactivity

It has been demonstrated that OECs exert immunomodulatory effects in OB under physiological condition (31). Moreover, the immunomodulatory role of bOEC transplantation after SCI has been highlighted in several studies (32). We therefore investigated both the *in vivo* and *in vitro* effects of bOEC transplantation on Mφ–Mi density and activation.

First, we counted Iba1+ cells in the rostral, caudal, and lesion-core regions of the spinal cord 14, 28, and 56 days post-SCI. This analysis revealed no significant differences between groups at any time point (Figure 5A and B). To further probe activation states, we assessed expression of CD86 and CD206, markers classically linked to pro-and anti-inflammatory phenotypes, respectively, and calculated CD86+/Iba1+ and CD206+/Iba1+ ratios in the same regions at 14, 28, and 56 days post-SCI (Figure 5C–H). Fourteen days after injury, only bOEC H transplantation lowered the CD86+/Iba1+ ratio in the rostral part of the spinal cord while raising the CD206+/Iba1+ ratio in both rostral and caudal segments relative to the SCI group. At this time point, bOEC I transplantation showed a similar trend, but the differences versus SCI were not significant (Figure 5C and D).

**Figure 5:**
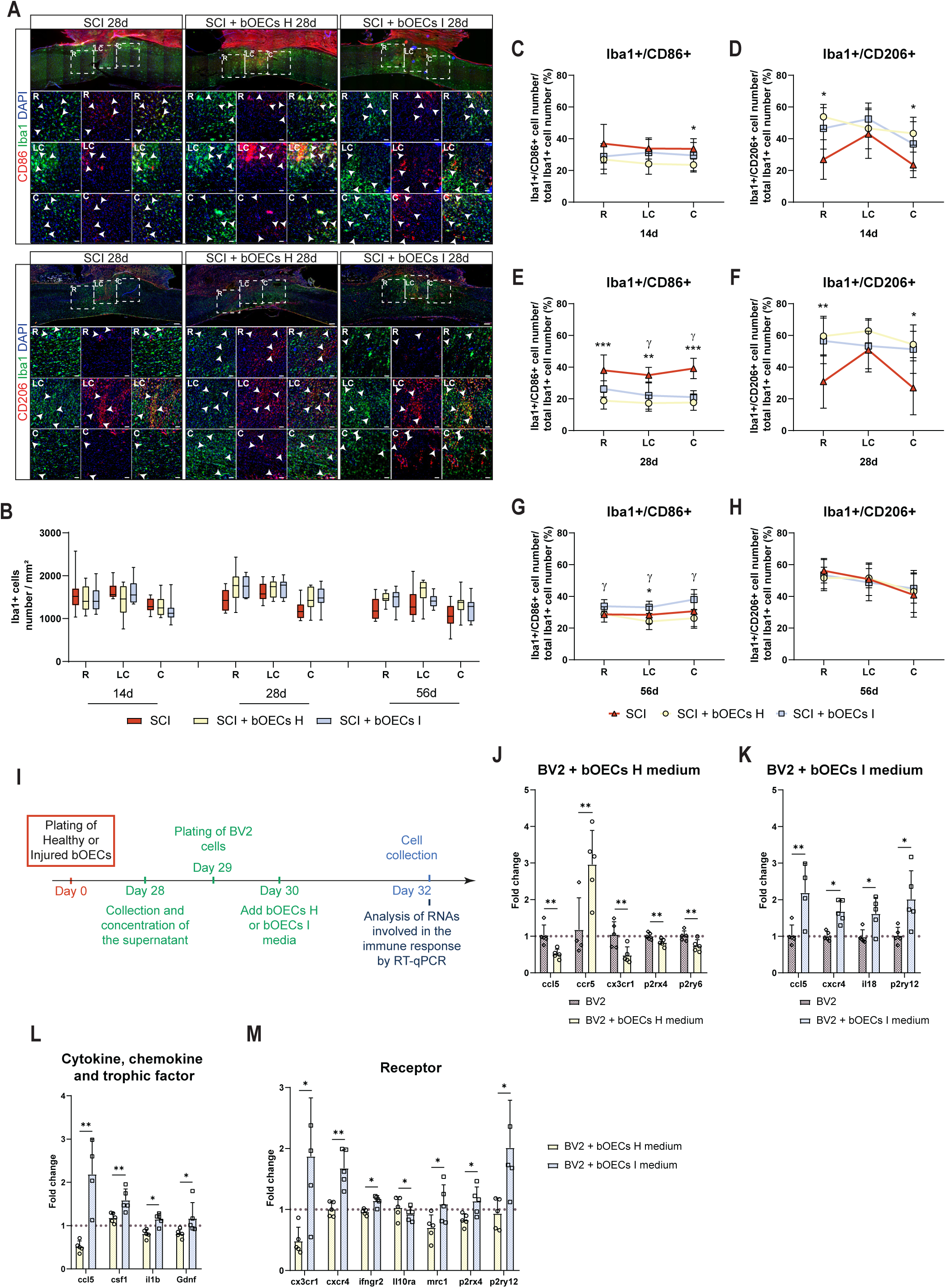
Transplantation of bOECs from healthy and injured mice alters microglia/macrophages polarization. **A-H**. Histological analysis of microglia/macrophage cell density and polarization. **A**. Representative images of sagittal spinal cord sections from SCI, SCI bOECs H and SCI bOECs I groups 28 days after SCI. Sections were stained with anti-Iba1, anti-CD86 antibodies and anti-CD206 antibodies. The square sections correspond to the rostral (R), lesion core (LC) and caudal (C) areas, defined for the quantification of Iba1+/CD86+ and Iba1+/CD206+ cells. Arrows indicate Iba1+/CD86+ microglial cells and Iba1+/CD206+ microglial cells. Scale bar = 250 µm (sagittal spinal cord sections) and 100 µm (Squares analysed in R, LC and C). **B.** The number of Iba1 positive microglia/macrophage cells was measured 14, 28 and 56 days after SCI. Microglia/macrophage polarization was analyzed by expression of pro-inflammatory marker CD86 (**C, E** and **G**) and anti-inflammatory marker CD206 (**D, F** and **H**) at 14d (**C** and **D**), 28d (**E** and **F**) and 56d (**G** and **H**) post SCI. **I-M**. Induction of BV2 cells polarization *in vitro* by bOECs conditioned medium (CM). **I**. Experimental design, healthy or injured bOECs were plated (day 0). Four weeks after plating, bOECs H and bOECs I CM were collected and concentrated (day 28). BV2 cells were plated (day 29) and the next day (day 30), bOECs H or bOECs I concentrated CM was added to the BV2 cell cultures. Two days after (day 32), BV2 cells were removed from the plates and analyzed by qRT-PCR. **J-M**. Histogram of mRNA expression of BV2 cell control and BV2 cells with bOECs H CM or bOECs I CM of cytokine, chemokine, trophic factor and receptor genes. Dashed lines correspond to mRNA expression from control BV2 cells cultures. Quantifications are expressed as average ± Min/Max (**B**) and average ± SD (**C**, **D**, **E**, **F, G**, **H**, **J**, **K**, **L** and **M**). N=5-10 (**B**), 8-10 (**C-F**) and 5-10 (**G** and **H**) animals per group. N=5 BV2 cell cultures (**J, K, L** and **M**) per group. Statistical analyses were based on Kruskal-Wallis (**B, C, D, E, F, G** and **H**) and Mann-Whitney (**J, K, L** and **M**) tests. * and γ = *P*< 0.05; ** = *P*< 0.01 and *** = *P*< 0.001.

By 28 days post-SCI, both transplanted groups modulate the Mφ–Mi activation profile; indeed, the CD86+/Iba1+ ratio was reduced in the lesion core and in the caudal segment in both groups and, additionally, in the rostral part of the spinal cord for the bOEC H group (Figure 5E). Conversely, the CD206+/Iba1+ ratio was increased in the rostral and caudal regions of both transplanted groups compared with SCI controls, reaching significance only for the bOEC H group (Figure 5F).

Interestingly, 56 days post-SCI, transplantation of bOECs I increased the ratio of CD86+/Iba1+ cells in the rostral, caudal, and lesion-core regions compared to the bOECs H group (Figure 5H). Moreover, the CD86+/Iba1+ ratio in the lesion core was also higher in bOECs I–transplanted mice than in the SCI group. No differences were detected for the CD206+/Iba1+ ratio among groups at this time point (Figure 5H).

To further investigate the influence of bOECs on microglial activation, we cultured BV2 cells, an immortalized mouse microglial cell line, either untreated or treated with CM from bOECs H or bOECs I, and performed qRT-PCR on BV2 transcripts from the three conditions (Figure 5I-M). Compared to control BV2 cultures, BV2 cells exposed to bOECs H CM down-regulated activation-stage genes (e.g., *p2ry6*, *p2rx4*) and pro-inflammatory genes (e.g., *ccl5*, *cx3cr1*), while up-regulating the pro-remyelinating gene *ccr5* (Figure 5J). By contrast, BV2 cells treated with bOECs I CM up-regulated pro-inflammatory genes such as *ccl5*, *cxcr4* and *il18* (Figure 5K). Direct comparison of the two CM conditions showed higher expression of the chemoattractant gene *ccl5* and inflammation-activating genes (e.g., *csf1*, *cx3cr1*) in the bOECs I CM cultures in comparison to the bOECs H CM cultures. Additionally, pro-inflammatory genes (e.g., *il1b*, *cxcr4*, *ifngr2*) were up-regulated, whereas the anti-inflammatory gene *il10ra* was down-regulated in BV2 cells treated with bOECs I CM relative to those treated with bOECs H CM (Figure 5L and M).

Taken together, these findings demonstrate that both bOEC transplants modulate Mφ–Mi gene expression and alter the profil of pro- and anti-inflammatory surface markers, but in distinct ways. Transplantation of bOECs H dampens Mφ–Mi activation and shifts cells away from a pro-inflammatory state both *in vitro and in vivo*. In contrast, bOECS I exert more heterogeneous effects, leading to either pro-inflammatory or anti-inflammatory modulation *in vivo* depending on the experimental time points, and a pro-inflammatory action *in vitro* on microglial cell cultures.

## DISCUSSION

The primary aim of our study was to determine whether the therapeutic capacity of cells harvested after SCI is impaired when they are later autografted to enhance spinal-cord regeneration. While the pathophysiological changes that occur within spinal tissue post-injury are well documented, few studies have examined the systemic cellular responses elicited by the trauma. However, it is now established that, in the days following SCI, circulating cytokines and chemokines can be detected in the serum of both rodents and humans (33). These factors are associated with peripheral immune-cell activation that affects multiple organs, including the spleen (34), kidneys, lungs (35), liver (33), adipose tissue (36), and brain (8). For instance, SCI has been linked to brain inflammation and cognitive deficits, particularly in regions such as the OBs, cortex, thalamus, and hippocampus. The spread of inflammation to the brain and peripheral tissues may therefore be a critical determinant of the efficacy of autologous cell therapy.

To investigate this question, we used a bOEC transplantation model. These cells, obtained from primary cultures of OBs, are well suited for transplantation because they are easy to cultivate, typically for just 2–4 weeks, and contain mainly two clearly identifiable cellular components: OECs and stromal cells. This culture approach also avoids purification steps that could intrinsically alter the cells, such as enzymatic treatment, antibody labeling, or cell -sorting procedures. In our study, we chose also to use cultures of OECs obtained from adult tissue samples in order to more closely mimic a clinical scenario, whereas in the literature, samples are often taken from very young animals (37,38). This sampling model is more representative of the human context but requires longer culture periods, in this case, four weeks.

Moreover, previous studies, ours and others, have shown that bOEC transplantation is more effective than grafts containing purified OECs or mucosal OECs (20,39), and the neuro-regenerative effects of bOEC transplantation are well documented. Following SCI, transplanted bOECs enhance functional recovery by modulating spinal-scar composition at both cellular and molecular levels. Analyses of bOEC-secreted factors demonstrate their ability to produce growth factors, extracellular-matrix-modulating enzymes, and molecules that facilitate axonal regrowth *in vitro* and *in vivo* (40). After transplantation, bOECs confer neuroprotection by limiting secondary injury and preserving axonal networks spared by the primary trauma (41). Through the release of metalloproteinases, they degrade axon-inhibitory ECM components, particularly CSPGs within the spinal scar, and stimulate the deposition of molecules that support axonal regrowth (42–44). They also modulate Mφ–Mi activation both directly, via cytokines and chemokines release, and indirectly, by preserving spinal-cord tissue, thereby dampening Mφ–Mi activation and creating an environment permissive for neuronal-network regeneration (45). Collectively, these effects enhance spinal-cord plasticity, promote axonal regrowth and remyelination, and stabilize neuronal circuits that support functional recovery (46).

Building on these findings, we assessed whether SCI alters the therapeutic potential of bOECs harvested 7 days after injury by comparing their transplantation outcomes (bOECs I) with those of bOECs from uninjured mice (bOECs H). In the early phase (14 days post-SCI), mice receiving bOECs I showed no significant functional improvement over SCI controls or bOECs H-treated animals; instead, their performance displayed high variability and only a modest, intermediate trend toward recovery (Figure 2B and C). By contrast, bOECs H transplantation produced clear functional gains.

This lack of functional benefit with bOECs I persisted into the late phase (56 days post-SCI), whereas bOECs H continued to confer superior recovery (Figure 3A and B). Sensory assessment with the Hargreaves test mirrored these results: at 14 days, bOECs H recipients withdrew the hind paw more rapidly than both SCI controls and bOECs I recipients (Figure 2E). At 56 days, sensitivity in bOECs H mice had normalized to SCI-control levels, whereas withdrawal latencies in the bOECs I group remained elevated (Figure 3C). Although this test is difficult in SCI animals (hind paws often rotate), the data nonetheless indicate better sensory recovery with bOECs H.

CatWalk analysis at 56 days, which combines motor and sensory readouts, confirmed these trends: bOECs H transplantation reduced the BOS (Figure 3E) and increased stride length (Figure 3F), whereas bOECs I produced smaller or absent improvements. Paw-pressure measurements did not differ significantly, yet they tended to be lower in bOECs H recipients, consistent with Hargreaves’ findings. Collectively, these results show that bOECs H outperform bOECs I in promoting early and sustained functional and sensory recovery after SCI.

To further explore why sensorimotor recovery differed between the grafted groups, we analysed spinal-scar composition. Transplantation of bOECs I did not modify the cellular composition of the scar relative to untreated SCI mice, whereas bOECs H transplantation reduced the fibrotic core and, consequently, the astrocyte-free zone at the early stage (Figure 2G and I). Likewise, only bOECs H decreased the demyelinated area (Figure 2J). By the late phase, 56 days after SCI, differences in scar cellularity tend to disappear: significant changes persisted only between the SCI and bOECs H groups (Figure 3J–M).

We also examined laminin, an ECM component that supports axonal growth. Fourteen days post-SCI, the laminin+/PDGFRβ+ area ratio was the same in both grafted groups and higher than in SCI mice (Figure 2H), indicating that bOECs I still conferred a molecular benefit despite their poorer cellular effect.

Because these outcomes might have reflected different graft compositions, we quantified OEC and stromal-cell proportions by flow cytometry. Cultures prepared 7 days after SCI contained approximately 30–40 % OECs and 60–70 % stromal cells, virtually identical to cultures from uninjured mice (Figure 1D) (20). Thus, the divergent *in-vivo* effects cannot be attributed to altered cell ratios but must arise from intrinsic differences between the cells. Supporting this conclusion, qRT-PCR showed that bOEC I cultures over-express genes coding for scar-forming molecules (Figure 1E). These genes drive stromal-cell and type-A pericyte proliferation and enhance astrocytic ECM deposition, ultimately promoting fibrotic-scar formation (6).

Functional and sensory recovery are not determined solely by scar formation; preservation or regeneration of neuronal networks after injury is also critical. Consistent with the observed up–regulation of genes involved in ECM modulation and axonal plasticity (Figure 1E), factors known to inhibit axonal regrowth (5), transplantation of bOECs I exerts only limited effects on axonal regeneration, yielding fewer vGlut1–positive synapses and no significant enhancement of axonal sprouting compared with bOECs H (Figure 4I–K and M).

Neuronal density and PNN represent additional determinants of post–injury functional recovery (28). Our data show that bOEC I transplantation fails to improve neuronal survival after SCI, whereas bOEC H transplantation increases neuronal numbers at 14- and 28-days post–injury (Figure 4B and C). WFA staining further reveals a higher PNN intensity in bOEC H recipients than in bOEC I ones at 14 days (Figure 4E). These findings support the notion that, in the early phase, an intact PNN in bOEC H–treated mice affords neuroprotection by limiting neuronal loss; as time progresses, the gradual reduction in PNN density may facilitate synaptic connectivity. Indeed, while the PNN shields neurons from an inflammatory milieu in the acute stage, its partial degradation is necessary later on to permit neuronal plasticity and the formation of new synapses (47).

The survival of the two grafted cell populations is identical and remains stable for at least one month after transplantation (Figure S1). Consequently, any influence of bOECs on spinal-cord repair is exerted mainly during the acute phase. At that stage, the immune system, particularly Mφ–Mi cells, is a major determinant of tissue preservation and regeneration; their numbers and polarization state strongly affect scar formation and axonal regrowth (48).

Our *in-vivo* analyses revealed no significant differences between the two grafted groups in Mφ– Mi density or overall polarization 14 and 28 days after SCI. Relative to SCI controls, however, both transplants modulate Mφ–Mi polarization toward an anti-inflammatory phenotype at early time points and curtailed the pro-inflammatory phenotype later on (Figure 5B–H). Because these measurements began on day 14, and early Mφ–Mi modulation is critical for repair, we next assessed the effects of bOECs H and I *in vitro*.

CM from bOECs I provoked marked up-regulation of cytokine, chemokine, trophic-factor, and receptor genes in BV2 microglial cultures (Figure 5K); most of these transcripts are linked to microglial activation and a pro-inflammatory state. By contrast, BV2 cells exposed to bOECs H CM showed a gene-expression profile close to that of untreated controls and even down-regulated several pro-inflammatory genes (Figure 5J). These observations align with the higher *il1β* and *tnf-α* expression detected in bOEC I primary cultures by qRT-PCR (Figure 1E). Together, the data indicate that bOECs I adopt a reactive state that drives Mφ–Mi toward a pro-inflammatory phenotype, whereas bOECs H exert a more immunomodulatory, anti-inflammatory influence.

Altogether, our findings demonstrate for the first time that the initial injury markedly influences the therapeutic efficacy of cells transplanted after SCI. Grafting bOECs I yield only modest functional recovery. This shortfall appears to stem from a reactive state in bOECs I that weakens their ability to modulate Mφ–Mi cells, leading to larger spinal scars and greater neuronal loss. The resulting increase in secondary damage, and the activation of less effective regenerative mechanisms, likely explains the intermediate benefit observed with bOEC I transplantation compared with bOECs H (Figure 6).

**Figure 6:** Schematic representation of our main results.

## Conclusion

Finally, our findings need to be viewed from a clinical standpoint. Autologous cell transplantation is being explored as a potential therapy for SCI, yet our data demonstrate that traumatic injury diminishes the neuro-regenerative capacity of cells harvested from injured animals. It is therefore reasonable to suspect that cells obtained from human patients with SCI would be similarly impaired, casting doubt on both the practicality and the therapeutic benefit of such autologous transplants.

## ACKNOWLEDGEMENTS

Behavioural studies and processing were conducted with the support of equipment from the Behavioural Analysis Platform SCAC (University of Rouen Normandy, France).

## USE OF AI

ChatGpt was used to correct the spelling, grammar and syntax of the sentences.

## Consent for publication

All authors have read the manuscript and indicated consent for publication

## Competing interests

The authors declare that they have no competing interests.

### AUTHOR CONTRIBUTIONS

Q.D. and N.G. conceptualized the project.

Q.D. and N.G. designed the experiments.

Q.D., M.B., P.N., L.M., C.R., A.R., A.H., A.B., F.S. and P.L. performed the experiments.

Q.D., M.B., L.M., A.H. and A.B. analyzed the results.

G.R. provided reagents, techniques and scientific input.

Q.D. and N.G. wrote the article.

---

**Figure S1:**
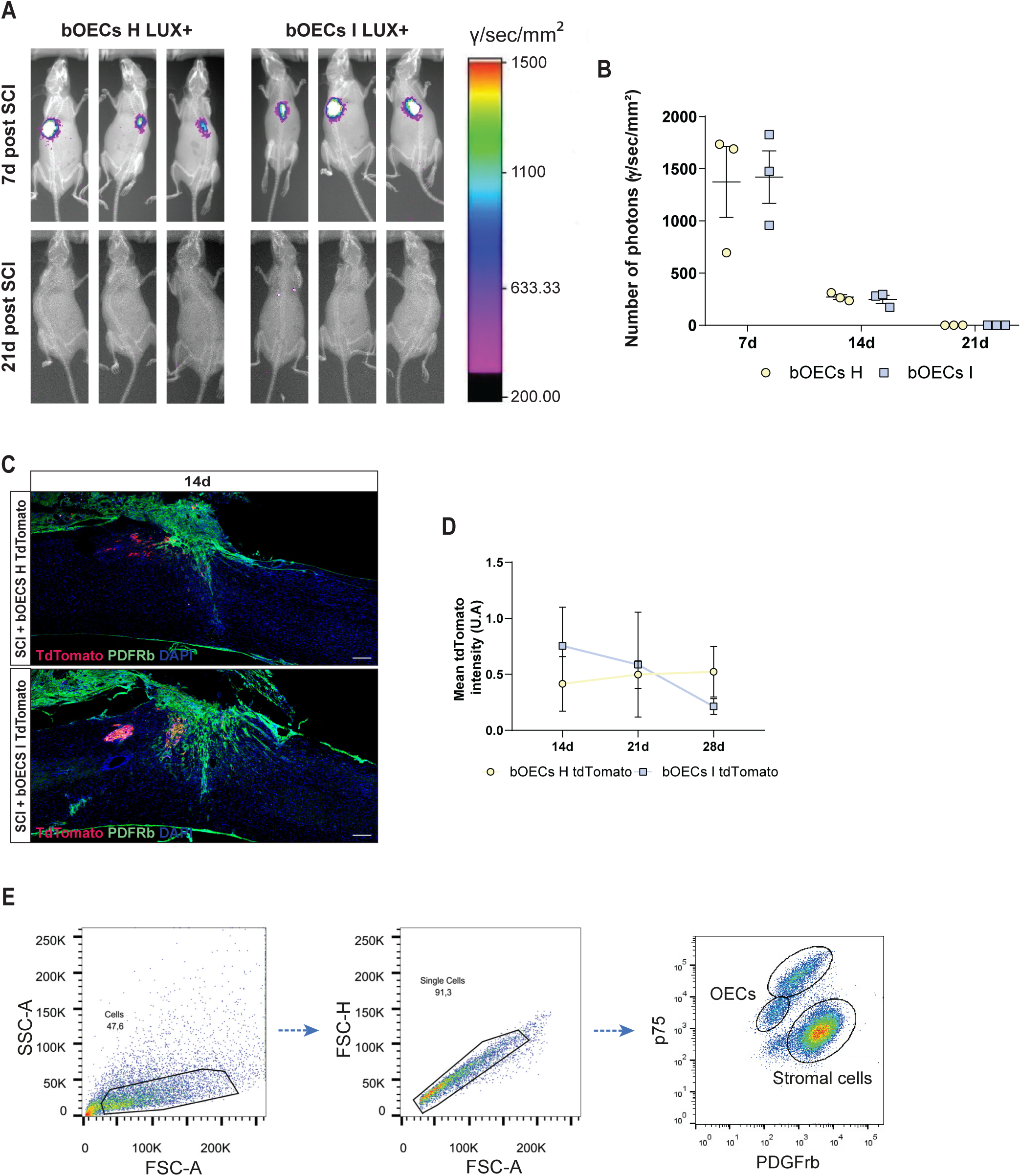
The survival of bOECs from injured mice is identical to bOECs’s survival from healthy mice after transplantation. **A**. Representative images of wild type mice transplanted with etiher LUX+/-bOEC H or LUX +/-bOEC I cultures 7 and 21 days after SCI. **B**. Quantification of luciferase+ signals (number of photons) 7, 14 and 21 days after SCI. **C**. Representative images of sagittal spinal cord sections from SCI bOECs H tdTomato and SCI bOECs I tdTomato groups 14, 21 and 28 days after SCI. Sections were stained with anti-PDGFrβ antibody. Dashed lines correspond to the injury site. Scale bar = 250 µm. **D**. Quantification of mean fluorescence intensity of the bOECs tdTomato H and I, 14, 21 and 28 days after SCI. **E.** Definition of gaits for cytometric characterization of bOECS primary cultures. Quantifications are expressed as average ± SD. N=3 (**B**) and 5 (**D**) animals per group. Statistical evaluations were based on Mann-Whitney tests.

**Table 1:**
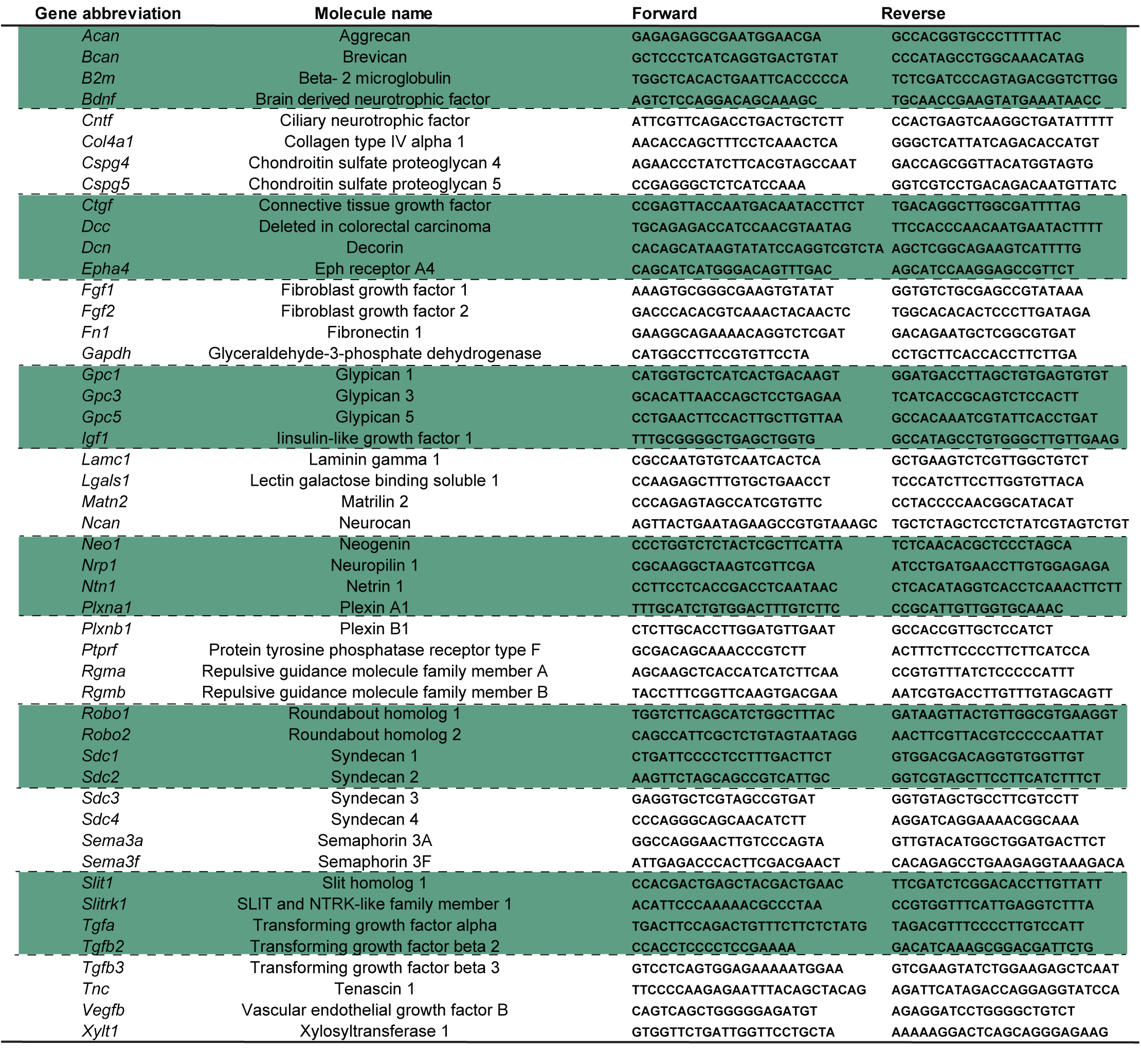
Table of genes for axonal growth inhibitory and axonal growth permissive molecules analyzed by qRT-PCR. This table lists the gene abbreviation, full name, forward and reverse primer sequences of 48 axon growth modulating molecules for which gene expression levels were analyzed in this study (5).

**Table 2:**
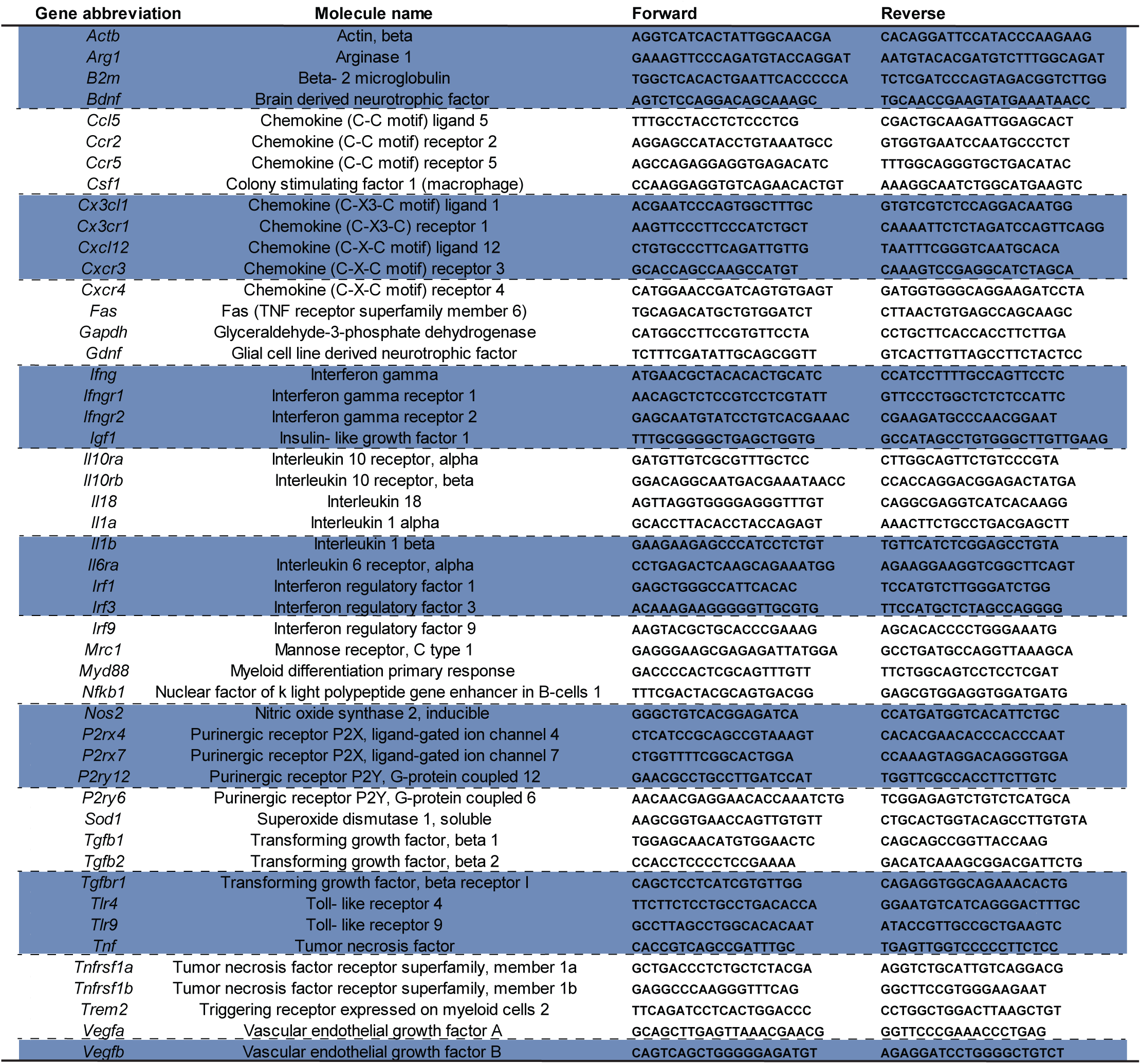
Table of genes for immunity and inflammatory molecules analyzed by qRT-PCR. This table lists the gene abbreviation, full name, forward and reverse primer sequences of 49 immune and inflammatory molecules for which gene expression levels were analyzed in this study (25).

## References

1. Liu Y, Yang X, He Z, Li J, Li Y, Wu Y, et al. Spinal cord injury: global burden from 1990 to 2019 and projections up to 2030 using Bayesian age-period-cohort analysis. Front Neurol. 2023 Dec 5;14:1304153.

2. Golestani A, Shobeiri P, Sadeghi-Naini M, Jazayeri SB, Maroufi SF, Ghodsi Z, et al. Epidemiology of Traumatic Spinal Cord Injury in Developing Countries from 2009 to 2020: A Systematic Review and Meta-Analysis. Neuroepidemiology. 2022;56(4):219–39.

3. Anjum A, Yazid MD, Fauzi Daud M, Idris J, Ng AMH, Selvi Naicker A, et al. Spinal Cord Injury: Pathophysiology, Multimolecular Interactions, and Underlying Recovery Mechanisms. Int J Mol Sci. 2020 Oct 13;21(20):7533.

4. Mortazavi MM, Verma K, Harmon OA, Griessenauer CJ, Adeeb N, Theodore N, et al. The microanatomy of spinal cord injury: a review. Clin Anat N Y N. 2015 Jan;28(1):27– 36.

5. Anderson MA, Burda JE, Ren Y, Ao Y, O’Shea TM, Kawaguchi R, et al. Astrocyte scar formation aids central nervous system axon regeneration. Nature. 2016 Apr;532(7598):195–200.

6. Göritz C, Dias DO, Tomilin N, Barbacid M, Shupliakov O, Frisén J. A pericyte origin of spinal cord scar tissue. Science. 2011 Jul 8;333(6039):238–42.

7. Sabelström H, Stenudd M, Réu P, Dias DO, Elfineh M, Zdunek S, et al. Resident neural stem cells restrict tissue damage and neuronal loss after spinal cord injury in mice. Science. 2013 Nov 1;342(6158):637–40.

8. Felix MS, Popa N, Djelloul M, Boucraut J, Gauthier P, Bauer S, et al. Alteration of Forebrain Neurogenesis after Cervical Spinal Cord Injury in the Adult Rat. Front Neurosci. 2012 Apr 9;6:19146.

9. Li Y, Ritzel RM, Khan N, Cao T, He J, Lei Z, et al. Delayed microglial depletion after spinal cord injury reduces chronic inflammation and neurodegeneration in the brain and improves neurological recovery in male mice. Theranostics. 2020 Sep 14;10(25):11376– 403.

10. Benabid AL, Costecalde T, Eliseyev A, Charvet G, Verney A, Karakas S, et al. An exoskeleton controlled by an epidural wireless brain-machine interface in a tetraplegic patient: a proof-of-concept demonstration. Lancet Neurol. 2019 Dec;18(12):1112–22.

11. Lorach H, Galvez A, Spagnolo V, Martel F, Karakas S, Intering N, et al. Walking naturall y after spinal cord injury using a brain–spine interface. Nature. 2023;618(7963):126–33.

12. Schneider MP, Sartori AM, Ineichen BV, Moors S, Engmann AK, Hofer AS, et al. Anti-Nogo-A Antibodies As a Potential Causal Therapy for Lower Urinary Tract Dysfunction after Spinal Cord Injury. J Neurosci Off J Soc Neurosci. 2019 May 22;39(21):4066–76.

13. Boido M, Garbossa D, Fontanella M, Ducati A, Vercelli A. Mesenchymal stem cell transplantation reduces glial cyst and improves functional outcome after spinal cord compression. World Neurosurg. 2014 Jan;81(1):183–90.

14. Yao S, Pang M, Wang Y, Wang X, Lin Y, Lv Y, et al. Mesenchymal stem cell attenuates spinal cord injury by inhibiting mitochondrial quality control-associated neuronal ferroptosis. Redox Biol. 2023 Sep 7;67:102871.

15. Pearse DD, Sanchez AR, Pereira FC, Andrade CM, Puzis R, Pressman Y, et al. Transplantation of Schwann cells and/or olfactory ensheathing glia into the contused spinal cord: Survival, migration, axon association, and functional recovery. Glia. 2007 juillet;55(9):976–1000.

16. Ursavas S, Darici H, Karaoz E. Olfactory ensheathing cells: Unique glial cells promising for treatments of spinal cord injury. J Neurosci Res. 2021 Jun;99(6):1579–97.

17. Assinck P, Duncan GJ, Hilton BJ, Plemel JR, Tetzlaff W. Cell transplantation therapy for spinal cord injury. Nat Neurosci. 2017 Apr 25;20(5):637–47.

18. Phelps PE, Ha SM, Khankan RR, Mekonnen MA, Juarez G, Ingraham Dixie KL, et al. Olfactory ensheathing cells from adult female rats are hybrid glia that promote neural repair. eLife. 2025 Apr 29;13.

19. Delarue Q, Mayeur A, Chalfouh C, Honoré A, Duclos C, Di Giovanni M, et al. Inhibition of ADAMTS-4 Expression in Olfactory Ensheathing Cells Enhances Recovery after Transplantation within Spinal Cord Injury. J Neurotrauma. 2019 Jul 2;37(3):507–16.

20. Gómez RM, Sánchez MY, Portela-Lomba M, Ghotme K, Barreto GE, Sierra J, et al. Cell therapy for spinal cord injury with olfactory ensheathing glia cells (OECs). Glia. 2018 Jul;66(7):1267–301.

21. Solstrand Dahlberg L, Becerra L, Borsook D, Linnman C. Brain changes after spinal cord injury, a quantitative meta-analysis and review. Neurosci Biobehav Rev. 2018 Jul;90:272– 93.

22. Wu J, Stoica BA, Luo T, Sabirzhanov B, Zhao Z, Guanciale K, et al. Isolated spinal cord contusion in rats induces chronic brain neuroinflammation, neurodegeneration, and cognitive impairment. Involvement of cell cycle activation. Cell Cycle Georget Tex. 2014;13(15):2446–58.

23. Takano M, Kawabata S, Shibata S, Yasuda A, Nori S, Tsuji O, et al. Enhanced Functional Recovery from Spinal Cord Injury in Aged Mice after Stem Cell Transplantation through HGF Induction. Stem Cell Rep. 2017 Mar 14;8(3):509–18.

24. Chort A, Alves S, Marinello M, Dufresnois B, Dornbierer JG, Tesson C, et al. Interferon β induces clearance of mutant ataxin 7 and improves locomotion in SCA7 knock-in mice. Brain J Neurol. 2013 Jun;136(Pt 6):1732–45.

25. Brifault C, Gras M, Liot D, May V, Vaudry D, Wurtz O. Delayed pituitary adenylate cyclase-activating polypeptide delivery after brain stroke improves functional recovery by inducing m2 microglia/macrophage polarization. Stroke. 2015 Feb;46(2):520–8.

26. Nakhjavan-Shahraki B, Yousefifard M, Rahimi-Movaghar V, Baikpour M, Nasirinezhad F, Safari S, et al. Transplantation of olfactory ensheathing cells on functional recovery and neuropathic pain after spinal cord injury; systematic review and meta-analysis. Sci Rep. 2018 Jan 10;8(1):325.

27. Chen Z, Fan H, Chen ZY, Jiang C, Feng MZ, Guo XY, et al. OECs Prevented Neuronal Cells from Apoptosis Partially Through Exosome-derived BDNF. J Mol Neurosci MN. 2022 Dec;72(12):2497–506.

28. Auer S, Schicht M, Hoffmann L, Budday S, Frischknecht R, Blümcke I, et al. The Role of Perineuronal Nets in Physiology and Disease: Insights from Recent Studies. Cells. 2025 Feb 20;14(5):321.

29. Anderson MA, O’Shea TM, Burda JE, Ao Y, Barlatey SL, Bernstein AM, et al. Required growth facilitators propel axon regeneration across complete spinal cord injury. Nature. 2018 Sep;561(7723):396–400.

30. Thornton MA, Mehta MD, Morad TT, Ingraham KL, Khankan RR, Griffis KG, et al. Evidence of axon connectivity across a spinal cord transection in rats treated with epidural stimulation and motor training combined with olfactory ensheathing cell transplantation. Exp Neurol. 2018 Nov;309:119–33.

31. Wright AA, Todorovic M, Murtaza M, St John JA, Ekberg JA. Macrophage migration inhibitory factor and its binding partner HTRA1 are expressed by olfactory ensheathing cells. Mol Cell Neurosci. 2020 Jan;102:103450.

32. Jiang C, Wang X, Jiang Y, Chen Z, Zhang Y, Hao D, et al. The Anti-inflammation Property of Olfactory Ensheathing Cells in Neural Regeneration After Spinal Cord Injury. Mol Neurobiol. 2022 Oct;59(10):6447–59.

33. Anthony DC, Couch Y. The systemic response to CNS injury. Exp Neurol. 2014 Aug 1;258:105–11.

34. Noble BT, Brennan FH, Popovich PG. The spleen as a neuroimmune interface after spinal cord injury. J Neuroimmunol. 2018 Aug 15;321:1–11.

35. Gris D, Hamilton EF, Weaver LC. The systemic inflammatory response after spinal cord injury damages lungs and kidneys. Exp Neurol. 2008 May 1;211(1):259–70.

36. Farkas GJ, Gorgey AS, Dolbow DR, Berg AS, Gater DR. The influence of level of spinal cord injury on adipose tissue and its relationship to inflammatory adipokines and cardiometabolic profiles. J Spinal Cord Med. 2018 Jul;41(4):407–15.

37. Pellitteri R, La Cognata V, Russo C, Patti A, Sanfilippo C. Protective Role of Eicosapentaenoic and Docosahexaenoic and Their N-Ethanolamide Derivatives in Olfactory Glial Cells Affected by Lipopolysaccharide-Induced Neuroinflammation. Mol Basel Switz. 2024 Oct 11;29(20):4821.

38. Tseng YT, Chen M, Lai R, Oieni F, Smyth G, Anoopkumar-Dukie S, et al. Liraglutide modulates olfactory ensheathing cell migration with activation of ERK and alteration of the extracellular matrix. Biomed Pharmacother Biomedecine Pharmacother. 2021 Sep;141:111819.

39. Mayeur A, Duclos C, Honoré A, Gauberti M, Drouot L, do Rego JC, et al. Potential of olfactory ensheathing cells from different sources for spinal cord repair. PloS One. 2013;8(4):e62860.

40. Delarue Q, Guérout N. Transplantation of Olfactory Ensheathing Cells: Properties and Therapeutic Effects after Transplantation into the Lesioned Nervous System. Neuroglia. 2022 Mar;3(1):1–22.

41. Khankan RR, Griffis KG, Haggerty-Skeans JR, Zhong H, Roy RR, Edgerton VR, et al. Olfactory Ensheathing Cell Transplantation after a Complete Spinal Cord Transection Mediates Neuroprotective and Immunomodulatory Mechanisms to Facilitate Regeneration. J Neurosci Off J Soc Neurosci. 2016 Jun 8;36(23):6269–86.

42. Gueye Y, Ferhat L, Sbai O, Bianco J, Ould-Yahoui A, Bernard A, et al. Trafficking and secretion of matrix metalloproteinase-2 in olfactory ensheathing glial cells: A role in cell migration? Glia. 2011 May;59(5):750–70.

43. Honoré A, Le Corre S, Derambure C, Normand R, Duclos C, Boyer O, et al. Isolation, characterization, and genetic profiling of subpopulations of olfactory ensheathing cells from the olfactory bulb. Glia. 2012 Mar;60(3):404–13.

44. Delarue Q, Robac A, Massardier R, Marie JP, Guérout N. Comparison of the effects of two therapeutic strategies based on olfactory ensheathing cell transplantation and repetitive magnetic stimulation after spinal cord injury in female mice. J Neurosci Res. 2021 Jul;99(7):1835–49.

45. Zhang L, Zhuang X, Kotitalo P, Keller T, Krzyczmonik A, Haaparanta-Solin M, et al. Intravenous transplantation of olfactory ensheathing cells reduces neuroinflammati on after spinal cord injury via interleukin-1 receptor antagonist. Theranostics. 2021;11(3):1147–61.

46. Radtke C, Akiyama Y, Brokaw J, Lankford KL, Wewetzer K, Fodor WL, et al. Remyelination of the nonhuman primate spinal cord by transplantation of H-transferase transgenic adult pig olfactory ensheathing cells. FASEB J Off Publ Fed Am Soc Exp Biol. 2004 Feb;18(2):335–7.

47. Grycz K, Głowacka A, Ji B, Krzywdzińska K, Charzyńska A, Czarkowska-Bauch J, et al. Regulation of perineuronal net components in the synaptic bouton vicinity on lumbar α-motoneurons in the rat after spinalization and locomotor training: New insights from spatio-temporal changes in gene, protein expression and WFA labeling. Exp Neurol. 2022 Aug;354:114098.

48. Zhou X, He X, Ren Y. Function of microglia and macrophages in secondary damage after spinal cord injury. Neural Regen Res. 2014 Oct 15;9(20):1787–95.

